# Post-acute immunological and behavioral sequelae in mice after Omicron infection

**DOI:** 10.1101/2023.06.05.543758

**Authors:** Tongcui Ma, Rahul K. Suryawanshi, Stephanie R. Miller, Katie K. Ly, Reuben Thomas, Natalie Elphick, Kailin Yin, Xiaoyu Luo, Nick Kaliss, Irene P Chen, Mauricio Montano, Bharath Sreekumar, Ludger Standker, Jan Mu□nch, F. Heath Damron, Jorge J. Palop, Melanie Ott, Nadia R. Roan

## Abstract

Progress in understanding long COVID and developing effective therapeutics is hampered in part by the lack of suitable animal models. Here we used ACE2-transgenic mice recovered from Omicron (BA.1) infection to test for pulmonary and behavioral post-acute sequelae. Through in-depth phenotyping by CyTOF, we demonstrate that naïve mice experiencing a first Omicron infection exhibit profound immune perturbations in the lung after resolving acute infection. This is not observed if mice were first vaccinated with spike-encoding mRNA. The protective effects of vaccination against post-acute sequelae were associated with a highly polyfunctional SARS-CoV-2-specific T cell response that was recalled upon BA.1 breakthrough infection but not seen with BA.1 infection alone. Without vaccination, the chemokine receptor CXCR4 was uniquely upregulated on multiple pulmonary immune subsets in the BA.1 convalescent mice, a process previously connected to severe COVID-19. Taking advantage of recent developments in machine learning and computer vision, we demonstrate that BA.1 convalescent mice exhibited spontaneous behavioral changes, emotional alterations, and cognitive-related deficits in context habituation. Collectively, our data identify immunological and behavioral post-acute sequelae after Omicron infection and uncover a protective effect of vaccination against post-acute pulmonary immune perturbations.

## INTRODUCTION

Long COVID (LC) is estimated to occur in at least 10% of individuals who recover from SARS-CoV-2 infection, and to affect at least 65 million individuals worldwide^1^. The factors driving LC pathogenesis are an active area of investigation, yet remain poorly defined, with SARS-CoV-2 persistence, herpes virus reactivation, autoimmunity, and overall immune dysregulation proposed as potential and non-mutually exclusive mechanisms^2-4^.

To date, studies deeply characterizing immune perturbations in LC have for the most part been limited to specimens from people, and such studies, while extremely relevant and important, harbor some limitations as compared to those using animal models. One challenge inherent to analysis of human specimens is that aside from blood, they are difficult to come by. As a result, to date few studies of LC in people (with some notable exceptions^5^) have analyzed tissues such as lungs, where SARS-CoV-2 takes an early foothold and can persist^6^, and where immune responses against the virus are most relevant to examine. Overall, much more is known about SARS-CoV-2-elicited immune responses in blood as compared to tissue compartments, although atlas studies from post-mortem tissues of people that succumed to acute severe COVID-19 have been informative in that regard^7^. Our studies have previously demonstrated in a severe mouse model of SARS-CoV-2 infection that extreme phenotypic changes, reflective of global T cell exhaustion, occur in the lungs of mice during the acute phase of severe COVID-19^8^. These studies took advantage of the ability of CyTOF – a high-parameter immunophenotyping tool – to deeply characterize total immune cell distributions and phenotypes, as well as the functional features of SARS-CoV-2-specific T cells. To date, however, such an in-depth immunophenotypic analysis has not been applied in the context of COVID-19 convalescence or vaccination. A second important challenge of human studies is that people are diverse, and all LC studies to date have been limited to some extent by the heterogeneity of the cohorts. Furthermore, controlled interventions are much more challenging in the human population than in mouse models.

One challenge to developing an animal model of LC is that the disease is complex, consisting of over 200 potential symptoms^9^, and is currently diagnosed through extensive interviewing of individuals that have recovered from COVID-19^10^. As it is not possible to implement a similar diagnostic approach in animals, *in vivo* models of LC have for the most part been limited to demonstrating persisting tissue pathologies and immune perturbations after the acute phase of SARS-CoV-2 infection^11-14^. In comparison, murine behavioral assays offer the opportunity to capture more nuanced features of LC. A murine model of LC exhibiting long-term memory deficits was recently developed through direct brain infusion of recombinant SARS-CoV-2 spike protein in wildtype Swiss Webster mice^15^, a mouse strain highly susceptible to encephalomyelitis. However, the long-term effects following intranasal SARS-CoV-2 infection have not yet been defined. Recent advances in machine learning and computer vision enable the assessment of spontaneous/naturalistic behavior and the identification of more subtle behavioral changes in mice. One such approach, termed variational animal motion embedding (VAME), has identified discrete behavioral motifs associated with amyloidosis and Alzheimer’s disease, that had escaped previous notice^16^. While VAME has not yet been used in the context of infection, we reasoned that it would provide a sensitive and unbiased means to detect the behavioral impact of SARS-CoV-2 infection in mice.

In this study, we used an animal model of mild COVID-19 where mice are exposed to Omicron BA.1, and analyzed these mice at convalescence and compared them to mice that were vaccinated in the absence or presence of infection. By implementing in-depth phenotyping by CyTOF, functional analysis of SARS-CoV-2-specific T cells, and VAME, we asked the following questions: 1) Do mice that have recovered from mild COVID-19 exhibit perturbations in immune compartments?; 2) To what extent does systemic mRNA vaccination elicit antigen-specific T cell responses in the lung and protect against any lingering perturbations caused by prior COVID-19?; and 3) Do mice that have recovered from mild COVID-19 exhibit behavioral changes? Overall, our findings provide an in-depth assessment of pulmonary SARS-CoV-2-specific immune responses in mice, and establish that mice that have recovered from mild COVID-19 serve as a valid model to study post-acute sequelae of COVID-19.

## RESULTS

### Perturbations in innate and B cells persist 3 weeks after BA.1 infection in unvaccinated but not vaccinated mice

We have recently shown that while the WA1 and Delta strains of SARS-CoV-2 are highly virulent and cause mortality in K18-hACE2 C57BL/6 mice, Omicron is mild and is cleared rapidly within about a week of infection, with all animals surviving^8^. We used this Omicron-based mild model of SARS-CoV-2 infection to assess for any persistent pulmonary immune cell changes that occur following intranasal (i.n.) inoculation with BA.1, as compared to exposure to SARS-CoV-2 antigen via systemic vaccination or vaccination followed by infection. We established four experimental groups, with 5 mice per group, as depicted in Fig. 1. Mice in Group 1 were never vaccinated nor infected (“Mock”). Mice in Group 2 were vaccinated intramuscularly (i.m.) with two doses of Moderna mRNA-1273 spaced 15 days apart (“Vac”). Mice in Group 3 were inoculated i.n. with 10^4^ PFU of BA.1 (“BA.1”). Mice in Group 4 were first vaccinated with mRNA-1273, rested for 30 days to enable development of immune memory, and then infected with BA.1 (“Vac+BA.1”). In all mice exposed to some form of antigen exposure (whether through vaccination or infection, Groups 2-4), sera were obtained 15 days following the final antigen exposure for assessment of antibody titers, and tissues were harvested 6 days later. The timepoint of tissue harvest was chosen to be 3 weeks following final antigen exposure to ensure that mice were no longer in the acute phase of infection, and to enable analysis of antigen-specific memory T cell responses. For tissue harvesting, lungs were digested as described in the Methods section. Single-cell suspensions of pulmonary cells were then subjected to CyTOF analysis at baseline, or following stimulation with a mixture of overlapping SARS-CoV-2 15-mer peptides and previously-validated and optimized SARS-CoV-2 epitopes in the C57BL/6 model^17^. Our CyTOF panel was specially designed to quantitate the major subsets of immune cells, to assess their homing potential, as well as to deeply phenotype and interrogate the effector functions of T cells (Table S1).

**Figure 1.**
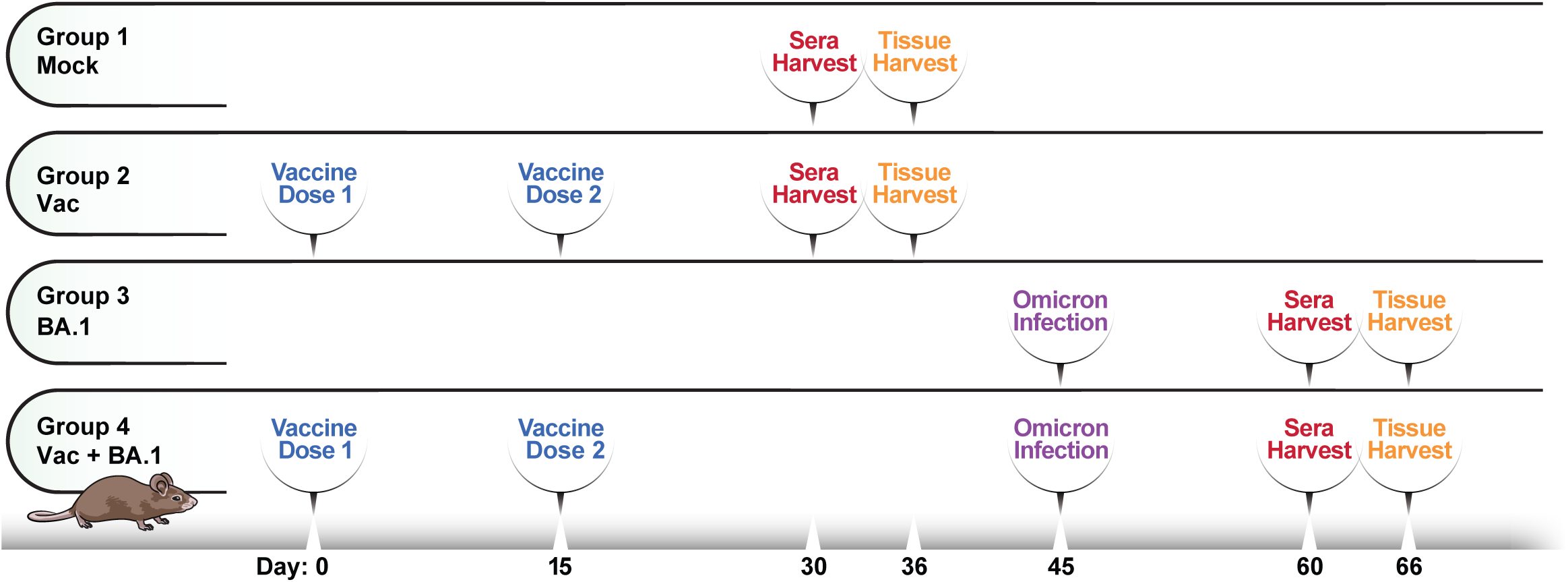
Schematic of experimental design for in-depth phenotyping of pulmonary immune cells. K18-hACE2 mice (n=5 / group) were divided into 4 experimental groups. Group 1 (Mock) consisted of mice that were never infected nor vaccinated. Group 2 consisted of mice that were vaccinated intramuscularly (i.m.) with mRNA-1273, and Group 3 consisted of mice that were infected intranasally (i.n.) with Omicron BA.1. Group 4 consisted of mice that were vaccinated, and then 30 days later infected i.n. with BA.1. In all cases, sera were obtained 15 days following the final antigen exposure (whether vaccination or infection), followed 6 days later by a harvest of lung specimens for immunophenotyping.

Comparison of subset distributions of immune cells from the lungs of the mice (manually gated as detailed in Fig. S1) revealed global changes, apparent both by tSNE visualization (Fig. 2A, S2) and by quantitation of manually-gated subsets (Fig. 2B). Notably, dendritic cells (DCs) (highlighted by the purple arrows in Fig. 2A) were significantly elevated in frequency after BA.1 infection, in a manner not observed in mice that were infected after vaccination (Fig. 2B). Granulocyte and NK cell frequencies also trended higher in the BA.1 convalescent mice relative to the other groups, while macrophage frequencies were similar between all groups (Fig. 2B). Although B cell frequencies were not significantly different between the BA.1 convalescent and mock-treated mice (Fig. 2C), global phenotypic alterations were apparent in the B cell compartment as evidenced by B cells from BA.1 convalescent mice occupying a unique region of tSNE space relative to mock-treated mice and the other experimental groups (Fig. 2A, dotted circle). The B cell perturbations in BA.1 convalescent mice were accompanied by a poor neutralizing SARS-CoV-2 antibody response (Fig. 2D). By contrast, all other treatment groups mounted antibody responses that neutralized not only the ancestral WA1 strain, but also the Alpha, Beta, Delta, and Omicron BA.1 and BA.2 subvariants (Fig. 2D). These results suggest that BA.1 infection, despite being mild, leads to lingering alterations in B cells and innate immune cells during convalescence, well after virus has been cleared.

**Figure 2.**
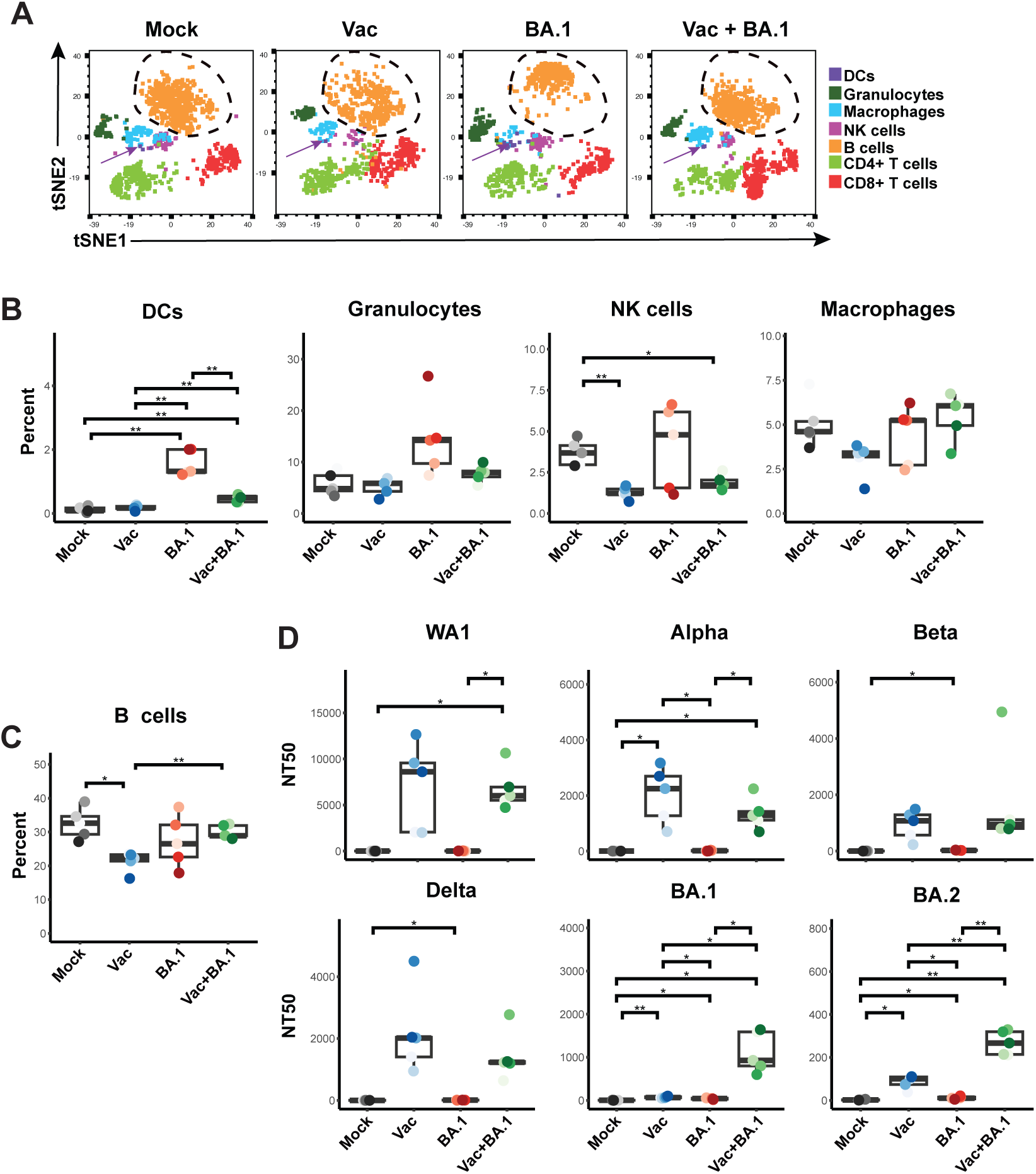
Innate and B cell perturbations persist during BA.1 convalescence. (**A**) Immune perturbations persist in the lungs of mice 3 weeks after BA.1 infection. Shown are total immune cells (CD45+) from mock-treated (Mock), mRNA-1273 vaccinated (Vac), BA.1 convalescent (BA.1), and mRNA-1273-vaccinated mice that then convalesced from BA.1 infection, (Vac+BA.1) (see details in Fig. 1). The indicated subsets of immune cells are color-coded according to the key on the right. Note that DCs (purple arrows) are increased in numbers and B cells (dotted circle) are phenotypically altered, in the BA.1 convalescent as compared to mock-treated mice, in a manner that is restored in the Vac+BA.1 group. Results are depicted as tSNE plots of representative mice from each experimental group, showing the same cell numbers for each group. Plots depicting cells from each individual mice analyzed in all groups are shown in *Fig. S2*. (**B**) Among innate immune cells in the lungs, DC frequencies are significantly increased in BA.1 convalescent mice, and granulocyte and NK cell frequencies trended higher. Macrophage frequencies were similar across experimental groups. Shown are box plots depicting the percentages of the indicated immune subset in mock, Vac, BA.1, and Vac+BA.1 mice. Percentages were calculated out of total immune cells. (**C**) Pulmonary B cell frequencies in BA.1 convalescent mice are not elevated relative to other experimental groups. Percentages were calculated out of total immune cells. (**D**) Neutralizing antibody responses are elevated in sera of both vaccinated experimental groups, but minimal in BA.1 convalescent mice. Neutralizing antibodies against each of the indicated SARS-CoV-2 isolates was measured using sera collected at timepoints detailed in Fig. 1. Results are presented as 50% neutralization titers (NT50s) against the respective SARS-CoV-2 isolates. *p<0.05, **p<0.01, as assessed using the Welch unpaired t-test and adjusted for multiple testing using the Benjamini-Hochberg for false discovery rate (FDR). In *panels B-D*, each datapoint corresponds to a different mouse.

### Minor perturbations occur in the T cell compartment of BA.1 convalescent mice

In contrast to the innate immune effector and B cells, no major alterations were observed in the frequencies and overall phenotypes of total CD4+ and CD8+ T cells of BA.1 convalescent mice relative to mock-treated mice (Fig. 2A, Fig. 3A, B). We did, however, observe some notable changes in BA.1 convalescent mice that were previously vaccinated (Vac+BA.1), relative to mock-treted mice. These included elevated frequencies of memory T cells and T resident memory cells (Trm), among both the CD4 and CD8 compartments, as well as CD4+ T follicular helper (Tfh) cells (Fig. 3A, B). These results suggest that vaccinated mice may be protected from the lingering innate and B cell perturbations associated with BA.1 infection (Fig. 2) through expansion of a robust recall T cell memory response elicited after BA.1 challenge.

**Figure 3.**
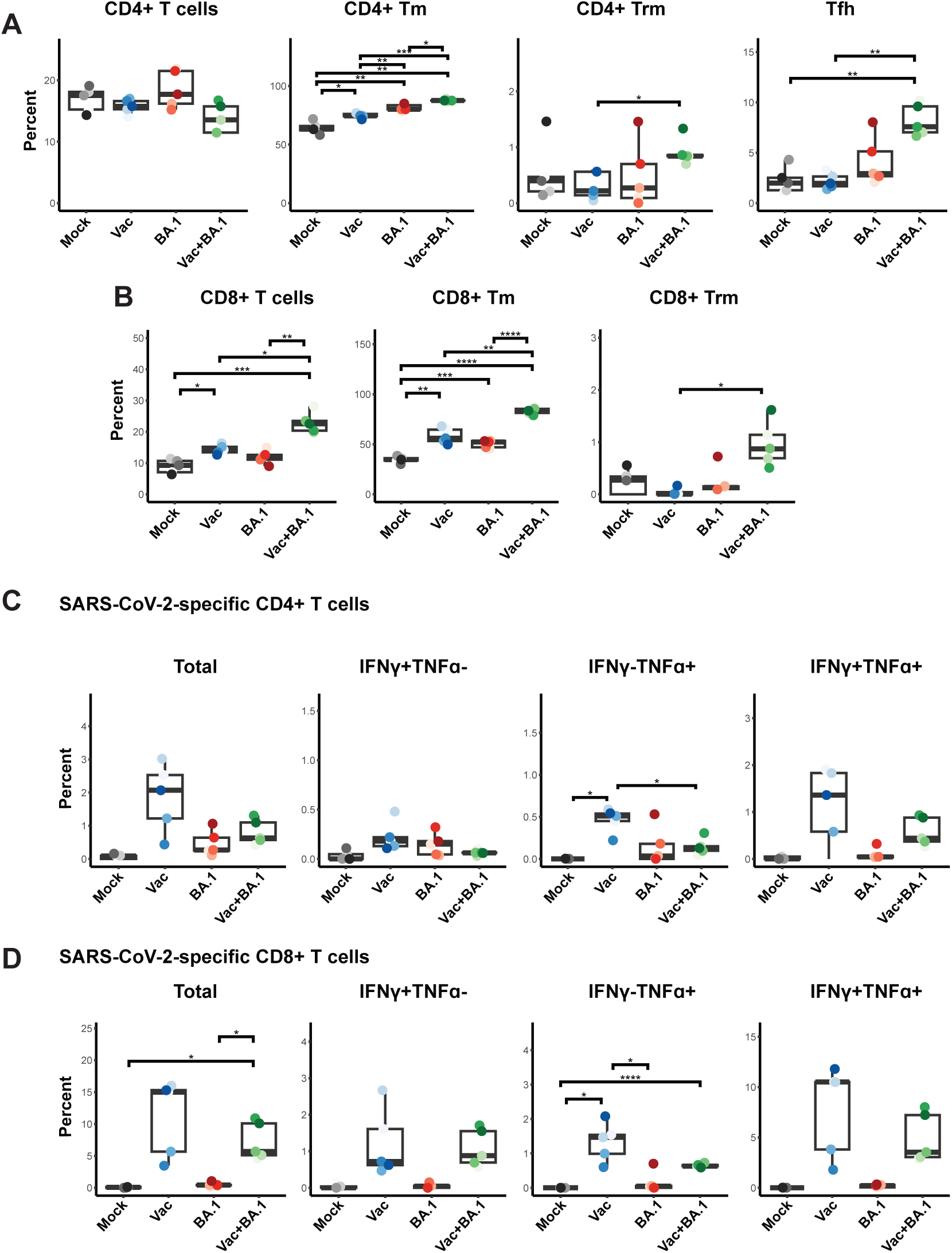
Pulmonary SARS-CoV-2-specific T cells are elicited by mRNA vaccination. (**A**) No major perturbations are observed among pulmonary CD4+ T cell subsets in BA.1 convalescent mice. Shown are the frequencies of pulmonary memory CD4+ T cells (CD4+ Tm), CD4+ T resident memory cells (CD4+ Trm), and T follicular helper cells (Tfh). The percentages of total CD4+ T cells were calculated out of total immune cells, while CD4+ Tm, CD4+ Trm, and Tfh percentages were calculated out of total CD4+ T cells. (**B**) No major perturbations are observed among pulmonary CD8+ T cell subsets in BA.1 convalescent mice. Shown are the frequencies of pulmonary CD8+ T cells, CD8+ Tm, and CD8+ Trm. The percentages of total CD8+ T cells were calculated out of total immune cells, while CD8+ Tm and CD8+ Trm percentages were calculated out of total CD8+ T cells. Note that the most prominent changes in T cell subsets were observed in the Vac+BA.1 group. (**C-D**) Robust induction of pulmonary SARS-CoV-2-specific CD4+ and CD8+ T cells producing IFNγ and/or TNFα in vaccinated but not BA.1 convalescent mice. Results are gated on live, singlet CD4+ (*C*) or CD8+ (*D*) T cells producing IFNγ and/or TNFα (Total), IFNγ only (IFNγ+ TNFα-), TNFα only (IFNγ-TNFα+), or both cytokines (IFNγ+TNFα+) in response to SARS-CoV-2 peptide stimulation. Each datapoint corresponds to a different mouse.

### mRNA vaccination elicits pulmonary SARS-CoV-2-specific T cells

To further interrogate the possibility that T cell immunity may play a role in protection from post-acute immune perturbations in the lung, we quantitated SARS-CoV-2-specific T cell responses from the lungs of these mice. Analysis of all effector markers in our CyTOF panel (IL2, IL5, IL6, IL10, IL17, IL21, IFNα, IFNγ, TNFα, perforin, granzyme B) revealed that IFNγ and TNFα could robustly and specifically identify populations of SARS-CoV-2-specific T cells by intracellular cytokine staining (Fig. S1). T cells inducing either of these two cytokines were therefore used to define SARS-CoV-2-specific T cells. SARS-CoV-2-specific CD4+ and CD8+ T cells trended highest in the Vac mice, and included those dually producing both IFNγ and TNFα (Fig. 3C, D). Interestingly, relative to Vac mice, the T cell response was lower in mice that were exposed to BA.1 after vaccination, despite the latter having been exposed to an additional dose of antigen in the lung. Of note, as the pulmonary immune environment of the Vac mice is one that is associated with subsequent protection against BA.1-associated post-acute immune perturbations (since Vac+BA.1 mice did not exhibit these perturbations), these results suggest that a robust SARS-CoV-2-specific T cell response in the lung may play a direct role in preventing post-acute sequelae of COVID-19. We therefore set out to more deeply characterize the features of these vaccine-elicited pulmonary SARS-CoV-2-specific T cells.

### Pulmonary SARS-CoV-2-specific T cells elicited by mRNA vaccination are polyfunctional exhibiting both cytokine and cytolytic effector functions

Assessing expression levels of all effector molecules on our CyTOF panel, we found that a subset of vaccine-elicited SARS-CoV-2-specific T cells producing IFNγ and/or TNFα also produced IL2 and granzyme B, while the remaining markers of effector function (IL5, IL6, IL10, IL17, IL21, IFNα, perforin) were not found to be produced in these cells (Fig. 4). Those eliciting both cytokine production (IFNγ, TNFα, or IL2) and cytolytic effector function (granzyme B expression) were observed among both the CD4+ (Fig. 4A) and CD8+ (Fig. 4B) T cells. Deeper interrogation of polyfunctionality by SPICE analysis^18^ revealed quadra-functional (TNFα+ IFNγ+ IL2+ granzyme+) SARS-CoV-2-specific T cells to be present among both the CD4 and CD8 compartments (Fig. 4C). Overall, almost half of the SARS-CoV-2-specific T cells produced at least 2 effectors (Fig. 4C). Together, these results suggest that a standard, systemically-administered 2-dose mRNA vaccination scheme elicits polyfunctional SARS-CoV-2-specific T cells that persist for at least 3 weeks in the lungs of mice.

**Figure 4.**
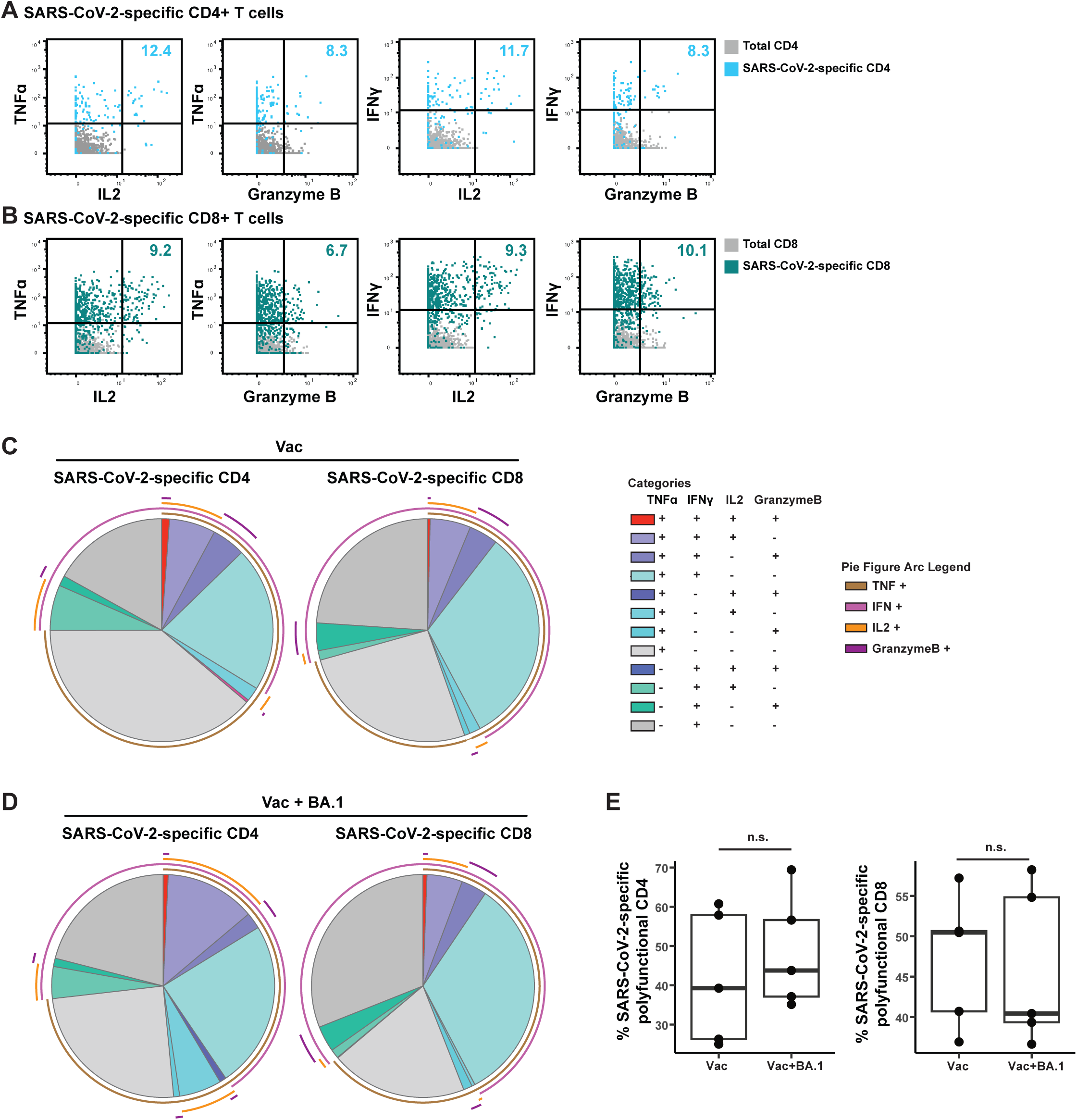
SARS-CoV-2-specific T cells in lungs of vaccinated mice are polyfunctional and can exhibit both cytokine and cytolytic effector functions. (**A, B**) Polyfunctional SARS-CoV-2-specific T cells are elicited in the lungs of mice vaccinated i.m. with mRNA-1253. Shown are 2D dot plots depicting the expression levels of TNFα, IFNγ, IL2, and granzyme B among SARS-CoV-2-specific CD4+ *(A)* or CD8+ *(B)* T cells. SARS-CoV-2-specific T cells are depicted in color, while total CD4+ or CD8+ T cells are depicted in grey. Numbers in the upper right of each plot indicate the percentages of SARS-CoV-2-specific CD4+ or CD8+ T cells producing the corresponding two effectors. (**C**) Polyfunctional SARS-CoV-2-specific T cells producing up to four effectors are elicited in the lungs of mice vaccinated i.m. with mRNA-1253. SPICE analysis was used to depict the proportions of SARS-CoV-2-specific CD4+ or CD8+ T cells producing various combinations of the effectors TNFα, IFNγ, IL2, and/or granzyme B. Cells producing all four effectors (red slice) were observed among both the CD4 and CD8 compartments. In total, 40% of SARS-CoV-2-specific CD4+ T cells and 47% of SARS-CoV-2-specific CD8+ T cells were polyfunctional. Mono-functional cell populations are depicted in shades of grey. (**D**) Polyfunctional SARS-CoV-2-specific T cells persist after recall response in the Vac+BA.1 mice. Data are presented as in *panel C*, with the identical color scheme. (**E**) Polyfunctionality of SARS-CoV-2-specific T cells is maintained before and after BA.1 infection. Shown are box plots of the percentages of SARS-CoV-2-specific CD4+ or CD8+ T cells expressing at least two of the effectors depicted in panels *C* and *D*.

### Comparison of the pre- vs. post-infection SARS-CoV-2-specific T cell response in vaccinated mice

In contrast to the Vac mice, Vac+BA.1 mice correspond to the setting where the T cells have completed their antiviral effector functions in response to breakthrough infection, restoring the lung to a state reminiscent of that in mock-treated mice. As such, in the context of vaccine-mediated protection against BA.1-induced perturbations, the Vac mice can be considered the “before” (pre-infection) state, and Vac+BA.1 mice the “after” (post-infection) state. We therefore next compared pulmonary SARS-CoV-2-specific T cells between these two groups of mice to assess T cell features preceding and succeeding breakthrough infection, in order to better understand the recall response that associates with restoration of lung homeostasis.

SPICE analysis revealed that the polyfunctionalities of both SARS-CoV-2-specific CD4+ and CD8+ T cells were maintained after BA.1 breakthrough infection (Fig. 4D, E). We reasoned that during a recall response, the phenotypes and differentiation states of SARS-CoV-2-specific T cells may also change. We therefore next implemented tSNE incorporating all T cell phenotyping markers on our CyTOF panel for data visualization. Of note, phenotypic analysis of SARS-CoV-2-specific T cells from mock-treated and BA.1 convalescent mice were not included as the frequencies of these cells in these groups were negligible/low (Fig. 3C, D). SARS-CoV-2-specific T cells from the Vac+BA.1 mice were phenotypically distinct from those of the Vac mice as demonstrated by their localizing in unique regions of tSNE space (Fig. 5A, B, S3). Clustering of the data by FlowSOM suggested different distributions of clusters in the two experimental groups (Fig. 5C, D), and this was particularly apparent for the CD8+ T cells. In particular, CD8+ T cell clusters 1, 4, and 5 were significantly more frequent in the breakthrough group, while Clusters 3, 6, and 10 were significantly less frequent (Fig. 5E, F). Interestingly, dual-functional (IFNγ+TNFα+) and mono-functional (IFNγ+TNFα-, IFNγ−TNFα+) SARS-CoV-2-specific T cells were all represented among the clusters over- and under-represented in the Vac+BA.1 mice, and these cytokines were major drivers separating these clusters from one another (Fig. 5G).

**Figure 5.**
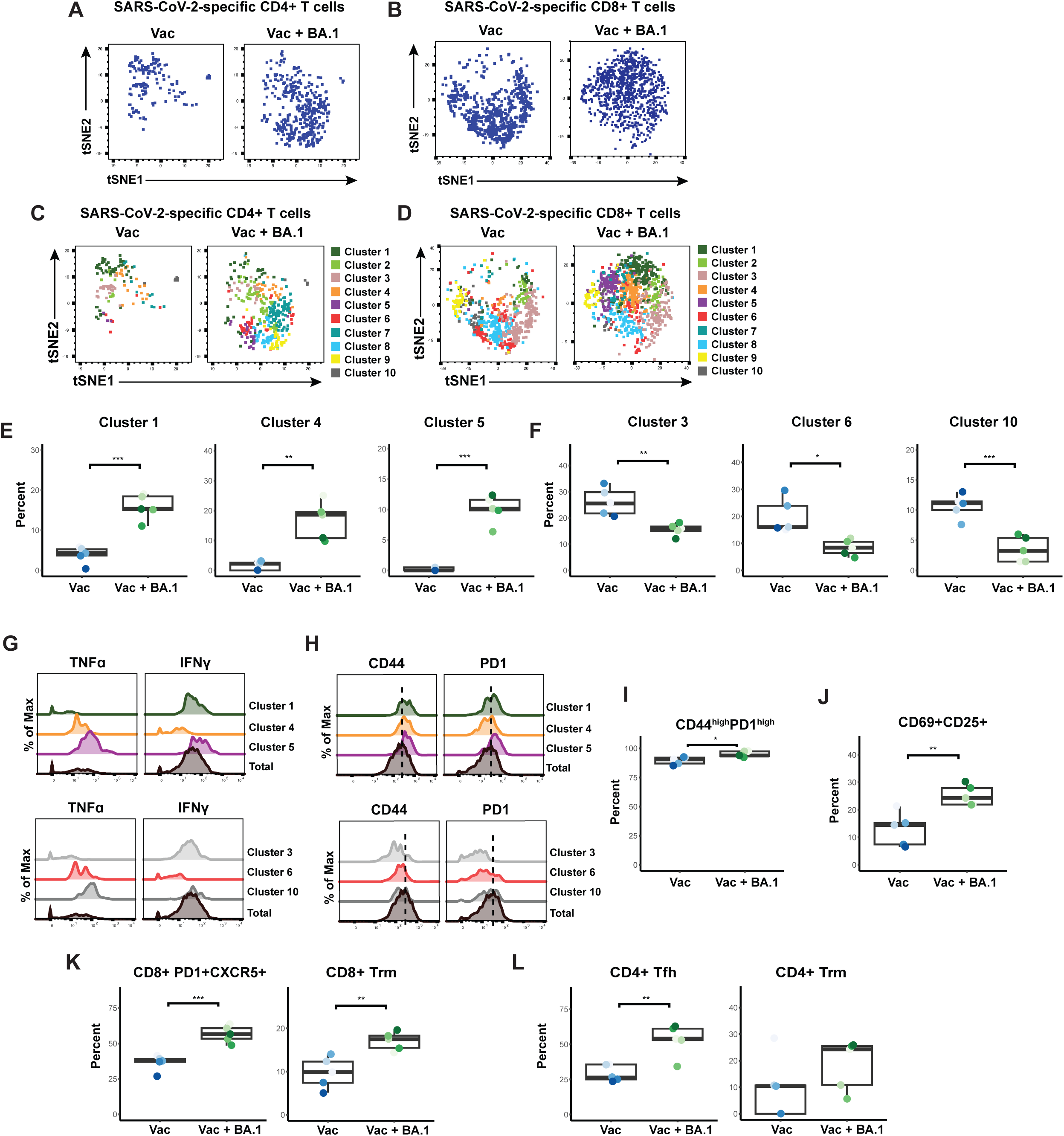
Pulmonary recall response to BA.1 breakthrough infection is characterized by increased T cell activation and enhanced follicular and T resident memory responses. (**A-B**) Pulmonary SARS-CoV-2-specific CD4+ (*A*) and CD8+ (*B*) T cells from Vac+BA.1 mice are phenotypically distinct from those of Vac mice, as suggested by different distribution of the cells in tSNE space. Shown are concatenated data from all mice within each gruop; data corresponding to each individual mouse are shown in Fig. S3. Note for depiction of the concatenated SARS-CoV-2-specific CD8+ T cells, the VA + BA.1 group was downsampled to match cell numbers in the Vac group to facilitate visualization. (**C-D**) Cluster distribution of SARS-CoV-2-specific CD4+ (*C*) and CD8+ (*D*) T cells differ in Vac+BA.1 mice. FlowSOM was used to cluster SARS-CoV-2-specific CD4+ and CD8+ T cells into 10 distinct clusters each. (**E**) Clusters 1, 4, and 5 of SARS-CoV-2-specific CD8+ T cells are significantly more abundant in Vac+BA.1 mice as compared to Vac mice. (**F**) Clusters 3, 6, and 10 of SARS-CoV-2-specific CD8+ T cells are significantly less abundant in Vac+BA.1 mice compared to Vac mice. (**G**) Clusters of SARS-CoV-2-specific CD8+ T cells differentially represented among Vac+BA.1 mice are driven by expression patterns of IFNγ and TNFα. Clusters 1 and 3 produce high levels of IFNγ without TNFα, Clusters 4 and 6 produce high levels of TNFα without IFNγ, and Clusters 5 and 10 produce high levels of both effector cytokines. (**H**) Expression levels of memory marker CD44 and activation marker PD1 are elevated on clusters of SARS-CoV-2-specific CD8+ T cells enriched in Vac+BA.1 mice. Dotted line corresponds to median expression intensity observed among total SARS-CoV-2-specific CD8+ T cells, highlighting the elevated (Clusters 1, 4, 5) or diminished (Clusters 3, 6, 10) expression of these antigens in the clusters. (**I**) SARS-CoV-2-specific CD8+ T cells from Vac+BA.1 harbor a higher proportion of CD44^high^PD1^high^ cells relative to those from Vac mice. (**J**) SARS-CoV-2-specific CD8+ T cells from Vac+BA.1 harbor a higher proportion of those dually expressing activation markers CD69 and CD25 as compared to those from Vac mice. (**K**) Among SARS-CoV-2-specific CD8+ T cells, those dually expressing PD1 and the follicle-homing marker CXCR5, and Trm cells, are significantly more abundant in Vac+BA.1 as compared to Vac mice. (**L**) Among SARS-CoV-2-specific CD4+ T cells, Tfh cells are significantly more abundant in Vac+BA.1 as compared to Vac mice. A similar trend is observed among SARS-CoV-2-specific CD4+ T cells of the Trm phenotype. *p<0.05, **p<0.01, ***p<0.001 as assessed using the Welch unpaired t-test and adjusted for multiple testing using the Benjamini-Hochberg for FDR.

Further interrogation of the clusters by expression levels of T cell phenotyping markers revealed that CD44 and PD1 were over-expressed in the clusters elevated in the Vac+BA.1 mice, with the opposite pattern observed in the clusters that were decreased in the Vac+BA.1 mice (Fig. 5H). We confirmed by manual gating that CD44^high^PD1^high^ cells were significantly enriched among SARS-CoV-2-specific CD8+ T cells from Vac+BA.1 mice (Fig. 5I). As PD1 is also an activation marker, we assessed whether the expression of other activation markers in our panel (CD69, CD25, CTLA4) were also elevated on SARS-CoV-2-specific CD8+ T cells in the Vac+BA.1 mice. While expression levels of CD69 and CD25 were significantly increased on the cells from Vac+BA.1 as compared to Vac mice, those of CTLA4 were not (not shown). Furthermore, CD69+CD25+ cells were more frequent among SARS-CoV-2-specific CD8+ T cells from the Vac+BA.1 as compared to the Vac group (Fig. 5J). PD1 and CD69 not only serve as activation markers but are used to define certain T cell subsets. In particular, PD1 co-expressed with CXCR5 defines T cells with important roles in follicular responses^19^, while CD69 is a marker of Trm. We found that among SARS-CoV-2-specific CD8+ T cells, those that were PD1+CXCR5+ or of the Trm phenotype were significantly enriched in the Vac+BA.1 group (Fig. 5K). Consistently, re-clustering of the SARS-CoV-2-specific CD8+ T cell data under conditions where cluster numbers were optimized by the Random Forests and a silhouette score-based assessment of clustering validity^20^ revealed two clusters, with the one significantly over-represented in Vac+BA.1 mice overexpressing PD1, CXCR5, CD69, and CD25 (Fig. S4). Interestingly, the CD4 equivalent of the subsets defined by these markers (Tfh, and CD4+ Trm) were also enriched among SARS-CoV-2-specific CD4+ T cells (Fig. 5L). Together, these findings demonstrate that activated follicle-homing and Trm cells are enriched in the mice that have experienced breakthrough infection.

### CXCR4 is a target for post-acute immune perturbations induced by BA.1 infection

Our data presented thus far support suggest that mild BA.1 infection induces post-acute perturbations in multiple immune subsets in the lung, but which prior mRNA vaccination protects against in a manner associated with a robust, polyfunctional pulmonary SARS-CoV-2-specific T cell response. These results are consistent with reports that vaccination decreases the risk of LC (in those that experienced breakthrough infection)^21,22^. However, as therapies are urgently needed for those who were infected prior to vaccination and subsequently developed LC, we wanted to assess whether our CyTOF datasets could reveal any potential targets, e.g., antigens that were differentially expressed in the BA.1 convalescent mice. Strikingly, one protein – CXCR4 – was highly over-expressed on multiple immune subsets in the lung in the BA.1 convalescent mice (Fig. 6, S5). CXCR4 is a chemokine receptor that is involved in homing of hematopoietic stem cells to the bone marrow, and can also direct immune cells to sites of inflammation. Its expression is associated with severe acute COVID-19^23^, and it has also been shown to be over-expressed on T cells from individuals with LC^4^. We found that CXCR4 was significantly over-expressed on granulocytes, macrophages, B cells, and CD4+ T cells in the context of BA.1 convalescence, and a similar trend was observed among DCs and NK cells. By contrast, in the context of breakthrough infection, CXCR4 profiles were restored to those indistinguishable from mock-treated mice (Fig. 6, S5). These observations together suggest that CXCR4 may be a promising therapeutic target for treatment of unvaccinated individuals who experienced post-acute sequelae of COVID-19.

**Figure 6.**
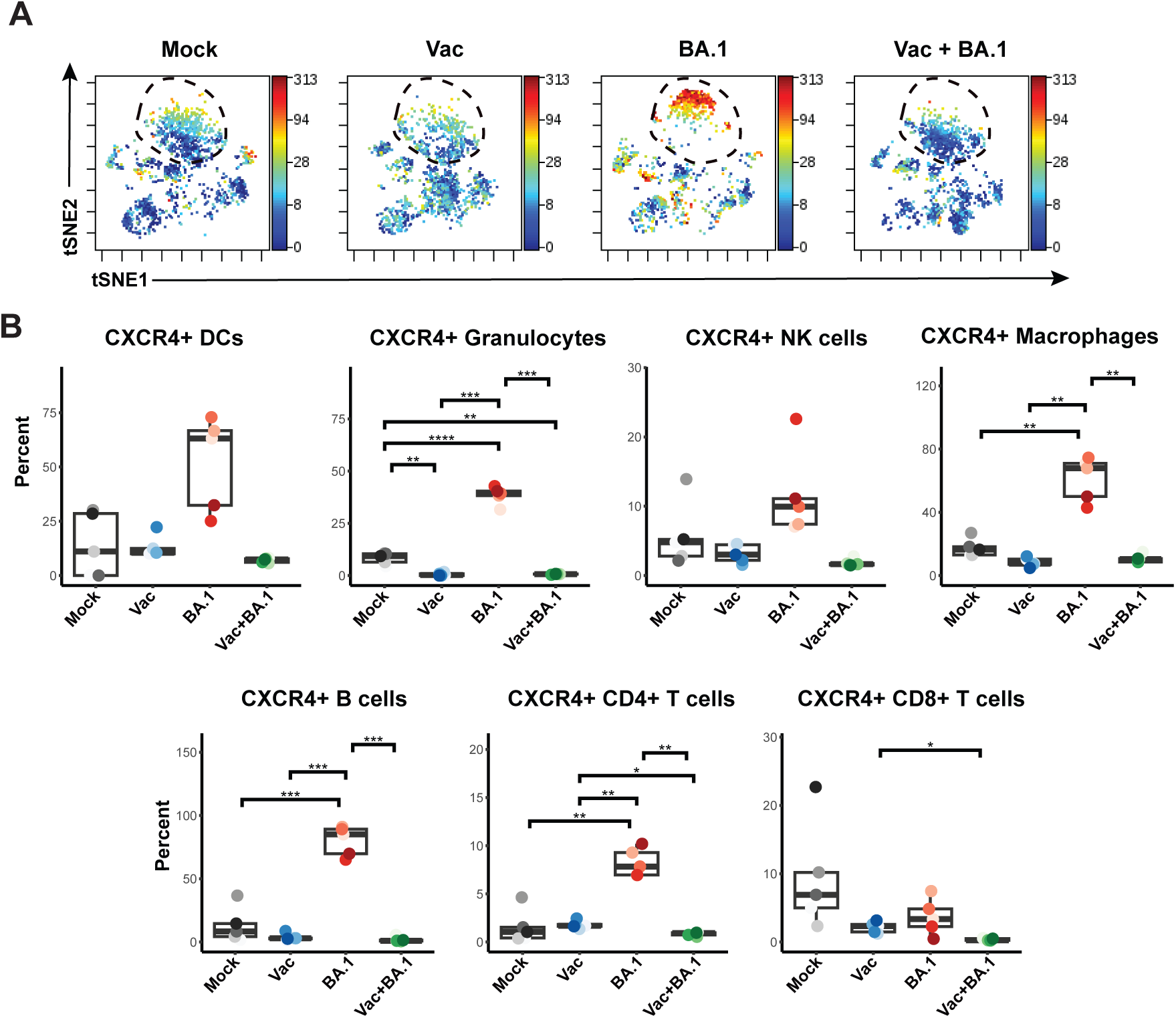
BA.1 convalescent mice, but not BA.1 convalescent mice previously vaccinated, harbor multiple subsets of pulmonary immune cells expressing high levels of CXCR4. (**A**) Pulmonary immune perturbations persisting 3 weeks after BA.1 infection is reflected by high expression levels of CXCR4, particularly among B cells (dotted line circles). tSNE maps are the same as those shown in Fig. 2A, but color-coded according to relative expression levels of CXCR4. (**B**) Multiple subsets of pulmonary immune cells from BA.1 convalescent mice express elevated levels of CXCR4. Lungs from BA.1 convalescent mice harbored significantly higher frequencies of CXCR4-expressing macrophages, granulocytes, B cells, and CD4+ T cells; positive trends were also observed among DCs and NK cells. *p<0.05, **p<0.01, ***p<0.001, ****p<0.0001 as assessed using the Welch unpaired t-test and adjusted for multiple testing using the Benjamini-Hochberg for FDR. Percentages were calculated out of total immune cells. Each datapoint corresponds to a different mouse.

### BA.1 convalescent mice exhibit post-acute sequelae as manifested by behavioral changes

Finally, we set out to determine whether the BA.1 convalescent mice with immunological perturbations exhibited behavioral changes. To enable in-depth computer-vision assessment of spontaneous/naturalistic behavior in an open arena, we established a novel double-containment setup in animal biosafety level 3 (ABSL3) conditions within a biosafety cabinet as detailed in Fig. S6. This system permits the capture of ventral images of spontaneous mouse behavior within an open arena. We selected a cohort of K18-hACE2 C57BL/6 mice, which were randomly assigned into two experimental groups: intranasal inoculation with vehicle (PBS) or with BA.1. Both groups underwent testing for 30 minutes prior to intranasal inoculation (Day 0 post-infection, or DPI 0; baseline) and again 3 weeks post-infection (DPI 21). Spontaneous behavior was quantified using standard and recently-developed machine learning approaches^16^.

At DPI 0, mice in the two groups did not exhibit significant differences in zone location (preferring periphery over center) nor in distance traveled (Figs. 7A, B), indicating comparable baseline conditions. Three weeks later, vehicle-treated mice maintained their preference for the periphery over center, whereas BA.1 convalescent mice no longer exhibited this preference, suggesting reduced anxiety or disinibition (Fig. 7A). BA.1 convalescent mice also traveled longer distances compared to vehicle-treated mice (Fig. 7B), suggesting a cognitive-related deficit in contextual habituation (the process by which mice exhibit less exploration when exposed to a familiar environment). Consistent with this, habituation scores (normalized distance traveled at DPI 21 relative to DPI 0) were significantly higher in BA.1 convalescent mice as compared to vehicle-treated mice (Fig. 7C). These results indicate that BA.1 convalescent mice exhibited impairments in cognitive (habituation impairment) and emotional (disinhibition) domains.

**Figure 7.**
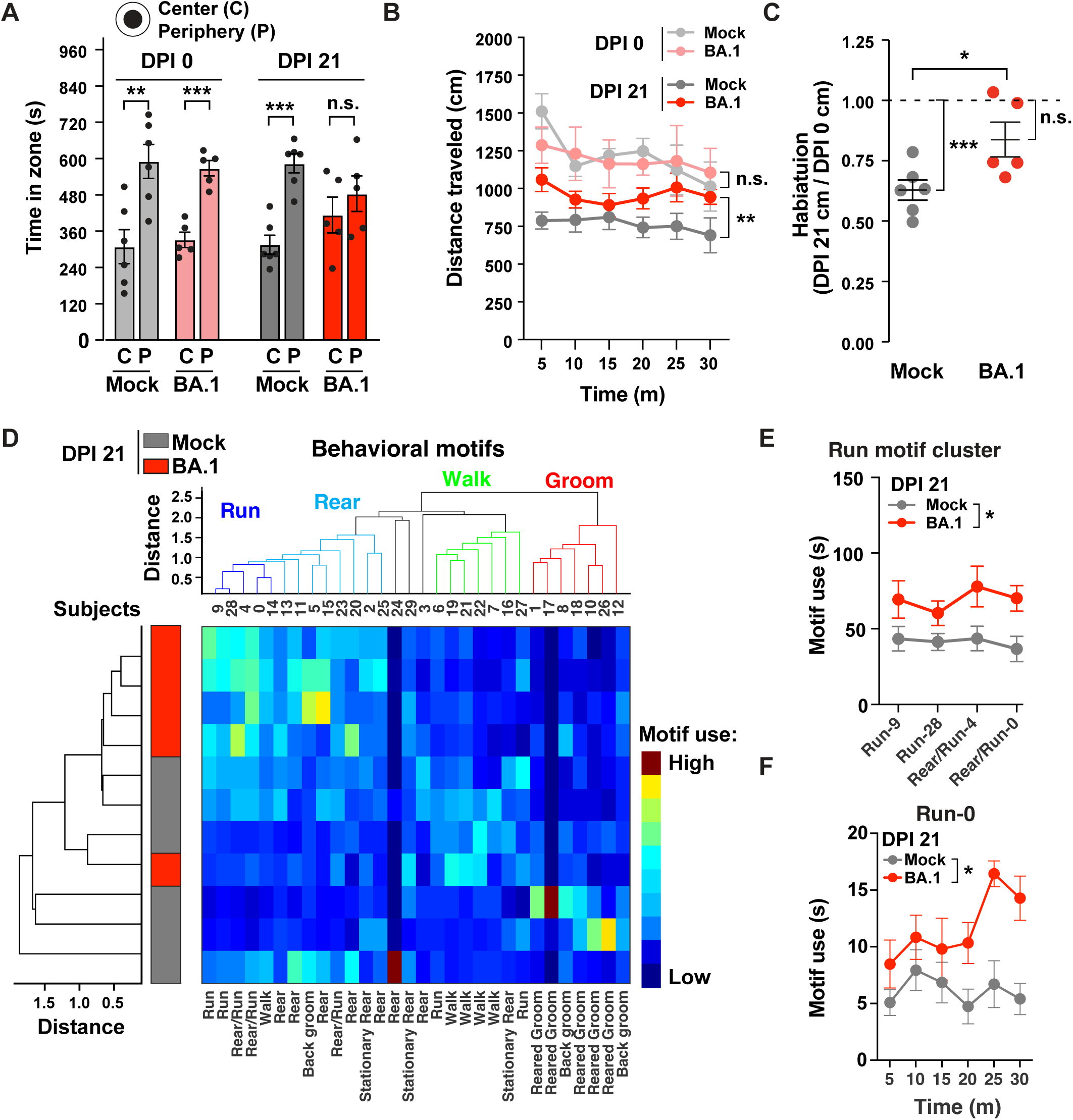
BA.1 convalescent mice exhibit behavioral alterations. Male mice were tested in an open arena for 30 minutes before (DPI 0) and 3 weeks after (DPI 21) intranasal inoculation with PBS vehicle (mock, n=6) or BA.1 (n = 5). (**A**) Time in center or periphery for the first 15 minutes of testing at DPI 0 and DPI 21. **p<0.01, ***p<0.001 by Student’s t-test. (**B**) Time-binned distance (5-minute bins) traveled in open field arena over 30 min at DPI 0 and DPI 21 in the two treatment groups. **p<0.001 by repeated one-way ANOVA test and Bonferroni tests for multiple comparisons. (**C**) Habituation scores (normalized total distance at DPI 21 relative to DPI 0). *p<0.05 by Student’s t-test. ***p<0.001 by one-sample Student’s t-test relative to 1. (**D**) Cluster analyses of motif use and subjects across all mice at DPI 21. Motif clusters (*top*: “run”, “rear”, “walk” and “groom”) and subject clusters (*left*) showed strong separation between vehicle and BA.1 convalescent mice. Heatmap represents motif use. (**E**) Total use of motifs within the “run” motif cluster (motifs 9, 28, 4 and 0) at DPI 21. *p<0.05 by two-way ANOVA. (**F**) Time-binned (5-minute bins) of “run” motif 0 use, over the 30 min of open arena test at DPI 21. *p<0.05 by two-way ANOVA.

To gain deeper insight into spontaneous behavioral alterations of BA.1 convalescent mice, we performed DeepLabCut (DLC) and VAME analyses^16,24^ at the DPI 21 timepoint. DLC^24^ was used to precisely track the location of 9 body parts (Fig. S6D, S7), and VAME^16^ was used to deconstruct mouse behavior into brief sequences of postural motifs. Cluster analyses of behavioral motif (motifs 0-29) use and subjects (mice 1-11) grouped motifs into multiple clusters, including “run”, “rear”, “walk” and “groom”, and revealed a strong separation between vehicle-treated and BA.1 convalescent mice (Fig. 7D). Overall, the motif clusters of “run” and “rear” were used more often by BA.1 convalescent mice, whereas those corresponding to “walk” and “groom” were used less often (Fig. 7D). The use of the “run” motif cluster (motifs 9, 28, 4 and 0) (Fig. 7E), as well as “run” motif 0 on its own (Fig. 7F), were particularly increased in the BA.1 convalescent mice. These results indicate that mice that had recovered from BA.1 infection exhibit significant and global changes in motif use, revealing profound changes in spontaneous behavior.

## DISCUSSION

In this study, we implemented high-parameter CyTOF phenotyping to characterize pulmonary immune cell changes in unvaccinated or vaccinated mice that had convalesced from mild BA.1 infection, and used standard and machine learning-based assessments of murine behavior to characterize post-acute sequelae of COVID-19 in mice. We demonstrate the ability of prior vaccination to protect against the persisting immune perturbations associated with prior infection, and characterize the phenotypes and effector functions of the vaccine-elicited T cell responses in the lung before and after SARS-CoV-2 exposure. We also report on CXCR4 as a potential new therapeutic target for those who suffer from LC, in particular those who were not vaccinated at the time of infection.

To our knowledge, this is first study which includes deep phenotypic analysis of the immunological features of mice that had convalesced from SARS-CoV-2 infection. We focused these analyses on the lung as it is the main site of viral replication during mild COVID-19, and pulmonary immune responses in the context of COVID-19 convalescence remain poorly characterized. We observed multiple immune perturbations in lungs of BA.1 convalescent mice, including an increase in the frequencies of DCs, granulocytes, and NK cells, as well as phenotypic alterations of macrophages and B cells characterized by high expression of CXCR4. We postulate that these changes reflect increased inflammation in the lung, which in the context of acute severe COVID-19 has been associated with the presence of CXCR4-expressing immune cells^23,25^. Strikingly, the immune perturbations observed 3 weeks after infection were absent when mice were vaccinated prior to infection. These observations suggest that vaccination can protect against post-acute sequelae of COVID-19, and are consistent with epidemiological observations in people^22^.

Our observation that systemic vaccination protected against immune perturbations in the lung prompted us to examine the extent to which the vaccine elicited a pulmonary immune response. We found that systemic administration of mRNA-1273 elicited a highly polyfunctional and robust T cell response in the lung within both the CD4 and CD8 compartments, that persisted for at least 3 weeks after vaccination. Effector functions of these cells included simultaneous production of IFNγ, TNFα, and IL2, which are cytokines associated with Th1 responses. By contrast, we did not find SARS-CoV-2-specific T cells producing the Th2 cytokine IL5, the Th17 cytokine IL17, or the immunosuppressive cytokine IL10. Intriguingly, SARS-CoV-2-specific T cells found in the lung included those that produced the cytolytic effector granzyme B, but as these cells did not express perforin, they may have a limited ability to mediate cytolysis of infected cells. This may serve to prevent excessive tissue damage in the lung, a lymphoid tissue which generally harbor few cytolytic T cells^26^. Upon BA.1 infection, the vaccine-elicited polyfunctional T cells were recalled back into action and further differentiated, particularly within the CD8 compartment. Overall, our findings point to an important role for vaccine-elicited T cell responses, including follicular and Trm responses, in protection against LC. This is in line with recent observations that people with LC exhibited increased frequencies of SARS-CoV-2-specific CD8+ T cells expressing multiple exhaustion markers, as it suggests that a sub-par CD8+ T cell response may be a mediator of the disease, potentially due to an inability to clear a persistent SARS-CoV-2 reservoir^4^.

There remains an unmet medical need for effective therapies for LC, particularly for those who were infected prior to vaccination. Our finding that CXCR4 was uniquely upregulated on multiple subsets of pulmonary immune cells in the context of COVID-19 convalescence points to CXCR4 antagonism as a potential approach for treatment. This is also supported by the observation that a subset of CXCR4-expressing cells from blood appears to be upregulated from cells of individuals with LC relative to those that recovered from COVID-19 without persisting symptoms^4^. Furthermore, a bioinformatics-based analysis of microarray PBMC data from 8 individuals with post-COVID inflammatory cardiomyopathy as compared to recovered COVID-19 convalescents found CXCR4 as a hub gene preferentially expressed in the former^27^. Interestingly, increased CXCR4 expression on immune cells is associated with aging^28^, and CXCR4 has also been suggested as a therapeutic target for severe/fatal acute COVID-19^23^. With regards to inhibitors, antagonism of CXCR4 by AMD3100 and potent peptide-based inhibitors has recently been reported to block lung inflammation and pulmonary infiltration of multiple immune cells (including eosinophils, T cells, B cells, and the neutrophil subset of granulocytes) in a CXCR4-driven model of asthma^29^. These inhibitors may serve as good candidates to test in future studies using our experimental mouse model of post-acute sequelae. More broadly, targeting of chemokine-receptor signaling cascades may also help ameliorate LC symptoms, given recent observations that auto-antibodies against multiple chemokines have been associated with protection against the disease^30^.

Finally, our data suggest that the machine learning approach VAME can be useful for establishing and characterizing LC, a complex disease for which suitable animal models are lacking. Despite the challenges of conducting behavioral testing in BSL3 conditions within a biosafety cabinet, our analyses demonstrated that significant behavioral changes persist for at least 3 weeks after COVID-19. This observation was not one we had expected, given that Omicron, unlike earlier strains like Delta or WA1, does not cause severe disease in mice^8^. Furthermore, compared to earlier strains, Omicron is associated with lower risk of LC, although the risk remains substantial^31^. However, as mild respiratory infection of mice with SARS-CoV-2 can lead to cellular dysregulation of the brain^32^, perhaps it was not completely unexpected that we could identify behavioral abnormalities in our system. The behavioral changes in the BA.1 convalescent mice which we observed through standard analyses and VAME included emotional alterations, cognitive-related deficits, and changes in spontaneous behavior. Changes in the emotional domain of the BA.1 convalescent mice were demonstrated by their no longer exhibiting preference for the periphery over the center in the open arena (disinhibition). Cognitive deficits in context habituation manifested as the diminished cognitive ability to recognize the same chamber the BA.1 convalescent mice were placed in 3 weeks prior, as reflected by increased exploration relative to control mice. Finally, spontaneous behavior alteration was demonstrated by global changes in motif use in the BA.1 convalescent mice. While fully acknowledging that LC is a complex disease that is difficult to recapitulate in a mouse model, we propose that the cognitive-related deficits in habituation may model cognitive symptoms or “brain fog” that is commonly reported in individuals with LC, and could be targeted for rescue by LC therapies in development. Future studies should expand the use of VAME to better define post-acute behavioral changes associated with COVID-19 convalescence. This should include examining sex differences (our study only examined male mice), since LC overall is more common in women^33^, as well as longer-term longitudinal studies. In addition, it will be of interest to perform VAME after SARS-CoV-2 infection in mouse models of neurodegenerative diseases such as Alzheimer’s, given the association of prior COVID-19 with worsening of neurological function^34^.

## METHODS

### SARS-CoV-2 virus preparation

Six million Vero-ACE2-TMPRSS2 cells cultured in T75 flasks were infected for 72 h at 37°C with 10 PFU of SARS-CoV-2 Omicron BA.1 or BA.2 (obtained as passage 1 stocks from the California Department of Health), or the pre-Omicron strains (obtained as passage 4-6) WA1 (USA-WA1/2020, from BEI), Alpha (B.1.1.7, from California Department of Health), Beta (B.1.351, from BEI), or Delta (B.1.617.2, from BEI). Supernatants of the cultures were used as viral stocks. The BA.1 and BA.2 supernatants were additionally concentrated using the Intact Virus Precipitation Reagent (Invitrogen). All viral stocks were stored at -80°C until use. The concentrations of viral stocks were determined using a plaque assay (as detailed further below) and quantified in plaque-forming units (PFUs). All viral infections were performed in the animal biosafety level 3 (BSL3) laboratory.

### Murine infection and vaccination

All protocols concerning animal use were approved by the Institutional Animal Care and Use Committee (IACUC) at the University of California, San Francisco (UCSF) and Gladstone Institutes (AN169239-01A). All animal studies were conducted in strict accordance with the National Institutes of Health (NIH) Guide for the Care and Use of Laboratory Animals. One week before the infection, mice were housed in an ABSL3 facility where the temperature (ranging from 65°F to 75°F) and humidity (maintained between 40% and 60%) were controlled. The mice experienced a 12-hour cycle of light and darkness, and had continuous access to water and standard laboratory rodent chow. For infections, 6-8 week-old K18-hACE2 male mice on the C57BL/6J background (bred in-house, originally obtained from Jackson Labs, #034860) were divided into four groups (n=5 per group) as depicted in Fig. 1. Two-dose mRNA-based COVID-19 vaccination was performed by inoculating mice twice intramuscularly (i.m.) with Moderna mRNA-1273 vaccine (50 μl of stock / mouse, equivalent to 10 μg or 1/10^th^ the human dose) spaced 15 days apart. Infections were established by exposing mice intranasally (i.n.) to BA.1 (10^4^ PFU in 50 μl PBS).

### Tissue processing and cell isolation

Murine lung tissues were harvested following transcardial perfusion. Harvested lungs were cut into smaller pieces, suspended in 100 μg/ml DNase I (Sigma-Aldrich) + 10 μg/ml Liberase TL (Sigma-Aldrich) in serum free-RPMI-1640 medium (Corning), and transferred to gentleMACS C-tubes (Miltenyi Biotec) before mounting on a GentleMACS dissociator (Miltenyi Biotec, program: m_Lung_01_02). The cells in the C-tube were pelleted by centrifugation, resuspended in CyFACS (metal contaminant-free PBS (Rockland) supplemented with 0.1% bovine serum albumin and 0.1% sodium azide) and sieved through a 70 μm strainer. Red blood cells (RBCs) were then lysed by resuspending cells in ammonium-chloride-potassium (ACK) lysis buffer (Thermo Fisher) for 2 min. The cells were then washed with CyFACS, and fixed for CyTOF analysis as detailed further below.

### Virus neutralization assay

Sera were serially diluted (1:30, 1:90, 1:270, 1:810, 1:2,430 and 1:7,290) in serum-free DMEM (Corning) to achieve final volumes of 50 μl for each dilution. The serum dilutions were separately mixed with 50 μl of SARS-CoV-2 WA1, Alpha, Beta, Delta, BA.1, BA.2 (50 PFU for each). The mixture was incubated at 37 °C for 30 min and then used in a plaque assay as described^8^. Plaques then were counted to calculate neutralization titers.

### Preparation of samples for CyTOF characterization of total immune cells and SARS-CoV-2-specific T cells

A total of 1-6×10^6^ cells per sample were exposed for 6 hours to media alone (R10 media, consisting of RPMI 1640 medium supplemented with 10% fetal bovine serum (FBS, VWR), 1% penicillin (GIBCO), and 1% streptomycin (GIBCO)), or to R10 containing 1 μM PepMix ancestral SARS-CoV-2 spike peptides (JPT Peptide Technologies) combined with 1 μM of a mix of previously reported optimal SARS-CoV-2 CD8+ T cell epitopes^17^ in order to identify SARS-CoV-2-specific T cells. Peptide stimulation was done in the presence of 3 μg/ml Brefeldin A Solution (eBioscience) to enable intracellular detection of induced cytokines. To enable subsequent normalization of specimens across multiple CyTOF runs, spleen cells from a single mouse were harvested and stimulated for 6 h with 16 nM PMA (Sigma-Aldrich) and 1 μM ionomycin (Sigma-Aldrich) in the presence of 3 μg/ml Brefeldin A Solution to generate a “bridge” sample run during each batch. Unstimulated, stimulated, and bridge sample cells were fixed for CyTOF analysis as described below.

### Fixation of cells for CyTOF analysis

Fixation of cells for CyTOF was performed similar to methods previously described^35,36^. Briefly, 1-6 million cells were washed once with contaminant-free PBS (Rockland) supplemented with 2 mM EDTA (PBS/EDTA), and then resuspended in 2 ml of the same buffer. The cells were then exposed for 60 seconds to an additional 2ml PBS/EDTA supplemented with 5 μM cisplatin, immediately after which cells were quenched with 10 ml CyFACS. The cells were then fixed for 10 min at room temperature with 2% PFA (Electron Microscopy Sciences) in PBS. After washing 3x with CyFACS, the cells were frozen at -80°C until CyTOF staining.

### CyTOF staining and data acquisition

Cells from multiple samples were combined together for staining using the cell-ID 20-Plex Pd Barcoding Kit (Standard BioTools) following the manufacturer’s protocol. In brief, 1-3 million cells were washed twice with Barcode Perm Buffer (Standard BioTools). After adding the appropriate barcode at a 1:90 (v/v) ratio, the cells were incubated for 30 min at room temperature, washed with 0.8 ml Maxpar^®^ Cell Staining buffer (Standard BioTools), followed by a second wash with 0.8 ml CyFACS. Barcoded samples were then combined and blocked for 15 min at 4°C with 100 μl of mouse and rat serum (both from Thermo Fisher) diluted at a 1:33 (v/v) ratio in CyFACS buffer. After washing twice with CyFACS, the cells were stained for 45 min at 4°C with a cocktail of CyTOF surface staining antibodies (Table S1) in a total volume of 100 μl / well. These CyTOF antibodies were purchased commercially (Standard BioTools), or conjugated in-house with X8 antibody-labeling kits (Standard BioTools) and stored at 4°C in Antibody Stabilizer (Boca Scientific) supplemented with 0.05% sodium azide. The stained cells were washed 3x with CyFACS buffer, and then fixed overnight at 4°C with 2% PFA in metal contaminant-free PBS. The next day, the cells were permeabilized by incubation with Fix/Perm buffer (eBioscience) for 30 min at 4°C, and then washed twice with Permeabilization Buffer (eBioscience). The cells were then blocked for 15 min at 4°C with 100 μl of mouse and rat serum diluted at a 1:5 (v/v) ratio in Permeabilization Buffer. After two washes with Permeabilization Buffer, the cells were stained for 45 min at 4°C with a cocktail of CyTOF intracellular staining antibodies (Table S1). After washing with CyFACS, the cells were incubated for 20 min at room temperature with PBS containing 250 nM Cell-ID^TM^ DNA Intercalator-Ir (Standard BioTools) and 2% PFA. Cells were then washed twice with CyFACS, and resuspended in 2% PFA diluted in PBS, and stored overnight in 4°C. The next day, immediately prior to data acquisition, the cells were washed sequentially with Maxpar^®^ Cell Staining Buffer, Maxpar^®^ PBS (Standard BioTools), and then Maxpar^®^ Cell Acquisition Solution (Standard BioTools). This was then followed by resuspending the cells at a concentration of 7 x 10^5^ / ml in EQ^TM^ calibration beads (Standard BioTools) diluted 1:10 in Maxpar^®^ Cell Acquisition Solution. Cells were acquired at a rate of 250-350 events/sec on a CyTOF2 instrument (Standard BioTools) at the UCSF Flow Cytometry Core.

### CyTOF data processing and normalization to bridge samples

Normalization to EQ^TM^ calibration beads and debarcoding of samples was performed using CyTOF software provided by Standard BioTools. The FCS files were generated and re-named with R (Version 4.2.1). Normalization of datasets against the bridge samples, which were included in each CyTOF run, was performed using CUHIMSR/CytofBatchAdjust^37^. All parameters of the experimental samples were adjusted using the 95th percentile batch normalization approach.

### Subset identification

Immune cells were defined as intact, live singlets expressing the pan-immune marker CD45. B cells were identified by co-expression of B220 and CD19, and T cells by co-expression of CD3 and TCRβ followed by CD4 and CD8 gating. After excluding B and T cells, NK cells were identified as those expressing NK1.1. Among the remaining NK1.1-cells, DCs were identified as those that were CD11c+. Then, among the remaining CD11c-cells, macrophages were identified as CD11b+Ly6G-cells, and granulocytes as CD11b+Ly6G+ cells. SARS-CoV-2-specific T cells were identified as those inducing IFNγ and/or TNFα upon SARS-CoV-2 peptide stimulation (Fig. S1). All manual gating was performed in FlowJo (BD). Memory T cells were defined as those expressing CD44. T follicular helper cells (Tfh) were defined as CD4+ T cells co-expressing PD1 and CXCR5, and resident memory T cells (Trm) were defined as T cells co-expressing CD69 and CD103.

### Box plot generation

The percentages of immune cells and T-cell subsets were exported from Flowjo, and the percentages of FlowSOM clusters were exported from Cytobank. The distribution of data across mice within each of the four (Mock, Vac, BA.1, and Vac+BA.1) or two (Vac and Vac+BA.1) experimental groups were visualized as box plots using the ggplot2 package^38^ in R. The lower, middle, and upper hinges of the box plots correspond to the 25th, 50th and 75th percentiles, respectively. The interquartile range (IQR) corresponds to the distance between the 25th and 75th percentiles. The upper whiskers indicates the largest value no further than 1.5X the IQR, while the lower whisker corresponds to the smallest value at most 1.5X the IQR. Data beyond the whiskers are outlier points.

### SARS-CoV-2-specific T cell characterization

For analysis of the polyfunctionality of SARS-CoV-2 specific T cells from vaccinated mice, we manually gated on CD4+ and CD8+ T cells specifically inducing TNFα, IFNγ, IL2, and/or granzyme B in response to SARS-CoV-2 peptide stimulation, and analyzed the data using Boolean gating in FlowJo. The frequencies of each population were exported and inputted into SPICE software (Version 6.1)^18^ for visualization of polyfunctionality.

### High-dimensional data analyses

FCS files corresponding to total immune cells and SARS-CoV-2-specific T cells were imported into Cytobank, which was then used for generation of tSNE plots. For mapping of different cell populations onto tSNE plots, the appropriate cellular subsets were defined based on standard and sequential two-dimensional dot plots, exported, and then pseudo-colored on the tSNE. FlowSOM was performed using default algorithm settings (clustering method: hierarchical consensus, metaclusters = 10, iterations = 10). Visualization of clusters by tSNE was performed in Flowjo. Additional clustering of SARS-CoV-2-specific CD8+ T cells was performed by optimizing the numbers of clusters by Random Forests^20^ together with a silhouette score-based assessment of clustering validity and cross-validation, as recently described^39^. To quantify cluster association in the Vac and Vac+BA.1 treatment groups, a generalized linear mixed model (GLMM) (implemented in the lme4^40^ package in R with the family argument set to the binomial probability distribution) was used to estimate the association between cluster membership and different treatment groups. The model consisted of cluster membership as a response variable, represented as the pair of the number of cells in and the number of cells not in the cluster under consideration. The treatment-specific log odds ratio of cluster membership association was estimated using the emmeans R package using the GLMM model fit.

### Behavioral Approaches

#### Naturalistic behavior in an open arena

For ventral video acquisition in the confines of a biosafety cabinet (Purifier Logic+ Class II, Type A2, Labconco), we created a custom behavioral setup and camera assembly (Fig. S6A). Naturalistic behavior was recorded in a clear acrylic circular chamber (12’’ diameter; 6’’ height) for 30 min during the day, avoiding the first and last two hours of the 7am-7pm light cycle. Two mice of the same treatment were recorded simultaneously in 2 independent chambers with visual screens between the mice. All recordings were made by the same researcher. Treatment groups were tested in alternating blocks throughout the day to minimize circadian effects as a potential confound. Chambers were cleaned with 70% ethanol between trials.

#### Videography

Machine learning video analysis requires high-quality ventral videos. Mice exploring an open chamber were recorded from below in RGB color at 30 frames per second with a GigE camera (acA1600-60gc, Basler), a 6 mm lens (UC Series, Edmund Optics), and Noldus Ethovision software. Four adjustable light bricks (Panel Go; Lume Cube) lit the chamber from below for homogeneous illumination of the paws and other body features. Videos were converted from .mpg to .mp4 format with FFmpeg before machine learning analysis.

#### Pose estimation

To track animal motion, we used DeepLabCut (DLC, version 2.1.8.2)^24^, a supervised machine learning tool which tracks animal posture by using deep convolutional neural networks. To create supervised annotations of key points, we extracted 10 frames from each of 31 videos and manually labeled 9 body parts (nose, front left paw, front right paw, back left paw, back right paw, belly, tailbase, midtail, tailtip) in each frame. The network (ResNet-50) was trained up to 460K iterations until the error converged (train error: 1.01 pixels, 0.75 mm; test error: 3.48 pixels, 2.57 mm) and precise virtual skeletons were estimated for all mice. Google Colaboratory (Colab Pro GPU) was used to execute network training and video analysis.

#### Behavioral segmentation and motif usage

The unsupervised behavioral segmentation tool VAME, was implemented previously^16^ to identify postural motions (motifs) from egocentric spatiotemporal information created from DLC-tracked key points. Motif analysis permits computation of kinematic variables that can be used to accurately identify complex ethologically-relevant behaviors. Briefly, mouse posture was egocentrically aligned in each frame to the belly and tailbase key points. Egocentrically-aligned coordinates and associated DLC confidence values were processed and used to train the VAME network, and 30 k-means clustered motifs were identified with a sliding time window of 533 milliseconds (16 frames). The midtail and tailtip coordinates were not used for VAME training, as these body parts reflect a time-lagged aspect of mouse motion. Motif speeds were defined as the median speed of the belly coordinate. Behavioral motif usage were defined as the total number of frames or seconds a mouse performed a given motif during the 30-min assay.

### Statistical Analysis

Welch unpaired t-tests were used for comparing the percentages of different immune cell types and antibody neutralization titers among experimental groups, and implemented in the package of rstatix^41^. The significance of the adjusted p-values was visualized on the box plots using the R package of ggprism^42^. Behavioral statistics were done with SPSS or Prism 9. As indicated in the figure legends, a one-sample t-test relative to 1 was used to assess habituation score changes, a two-sample t-test was used to assess location preference in the open arena, and repeated measures one-way ANOVA was used to assess longidutinal changes in distance or motif use. Two-tailed p-values less than 0.05 after adjustment were considered significant.

### Data Availability

Raw CyTOF datasets are available for download through the following link in the Dryad public repository database: https://doi.org/10.7272/Q62Z13RT

---

**Table S1.**
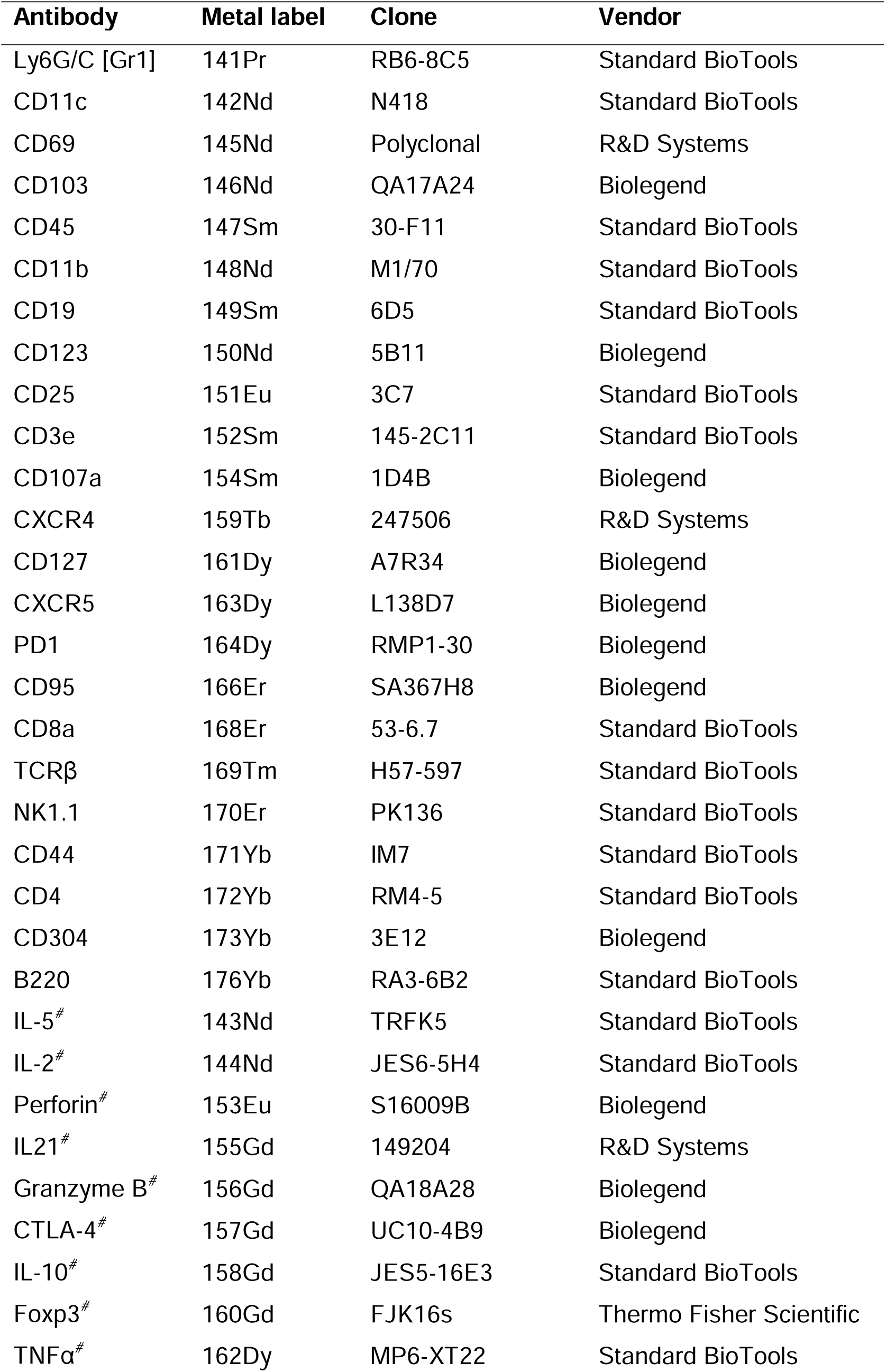
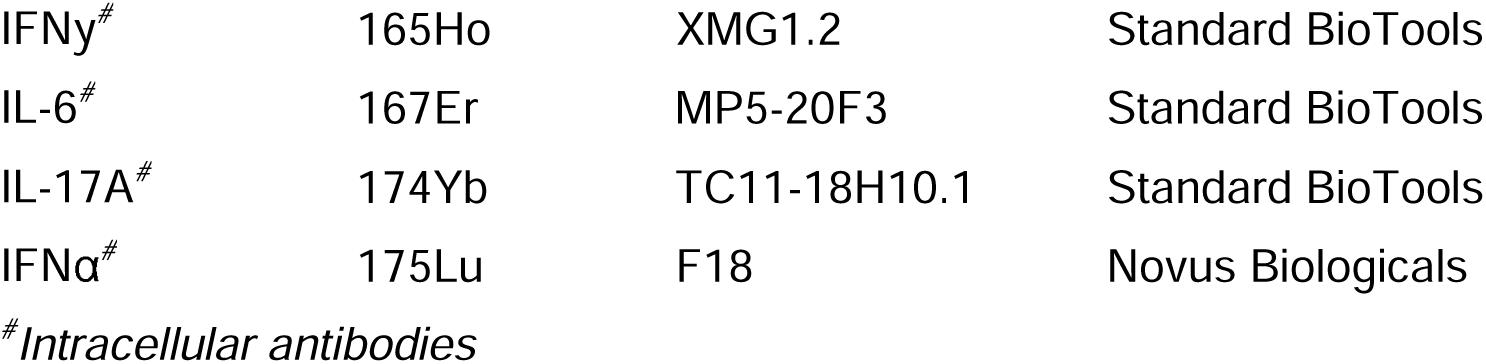
Mouse CyTOF antibodies.

| **Antibody** | **Metal label** | **Clone** | **Vendor** |
| --- | --- | --- | --- |
| Ly6G/C [Gr1] | 141Pr | RB6-8C5 | Standard BioTools |
| CD11c | 142Nd | N418 | Standard BioTools |
| CD69 | 145Nd | Polyclonal | R&D Systems |
| CD103 | 146Nd | QA17A24 | Biolegend |
| CD45 | 147Sm | 30-F11 | Standard BioTools |
| CD11b | 148Nd | M1/70 | Standard BioTools |
| CD19 | 149Sm | 6D5 | Standard BioTools |
| CD123 | 150Nd | 5B11 | Biolegend |
| CD25 | 151Eu | 3C7 | Standard BioTools |
| CD3e | 152Sm | 145-2C11 | Standard BioTools |
| CD107a | 154Sm | 1D4B | Biolegend |
| CXCR4 | 159Tb | 247506 | R&D Systems |
| CD127 | 161Dy | A7R34 | Biolegend |
| CXCR5 | 163Dy | L138D7 | Biolegend |
| PD1 | 164Dy | RMP1-30 | Biolegend |
| CD95 | 166Er | SA367H8 | Biolegend |
| CD8a | 168Er | 53-6.7 | Standard BioTools |
| TCRβ | 169Tm | H57-597 | Standard BioTools |
| NK1.1 | 170Er | PK136 | Standard BioTools |
| CD44 | 171Yb | IM7 | Standard BioTools |
| CD4 | 172Yb | RM4-5 | Standard BioTools |
| CD304 | 173Yb | 3E12 | Biolegend |
| B220 | 176Yb | RA3-6B2 | Standard BioTools |
| IL-5<sup>#</sup> | 143Nd | TRFK5 | Standard BioTools |
| IL-2<sup>#</sup> | 144Nd | JES6-5H4 | Standard BioTools |
| Perforin<sup>#</sup> | 153Eu | S16009B | Biolegend |
| IL21<sup>#</sup> | 155Gd | 149204 | R&D Systems |
| Granzyme B<sup>#</sup> | 156Gd | QA18A28 | Biolegend |
| CTLA-4<sup>#</sup> | 157Gd | UC10-4B9 | Biolegend |
| IL-10<sup>#</sup> | 158Gd | JES5-16E3 | Standard BioTools |
| Foxp3<sup>#</sup> | 160Gd | FJK16s | Thermo Fisher Scientific |
| TNFα<sup>#</sup> | 162Dy | MP6-XT22 | Standard BioTools |
| IFNγ<sup>#</sup> | 165Ho | XMG1.2 | Standard BioTools |
| IL-6<sup>#</sup> | 167Er | MP5-20F3 | Standard BioTools |
| IL-17A<sup>#</sup> | 174Yb | TC11-18H10.1 | Standard BioTools |
| IFNα<sup>#</sup> | 175Lu | F18 | Novus Biologicals |

<sup>#</sup>*Intracellular antibodies*

## FIGURE LEGENDS

**Figure S1.**
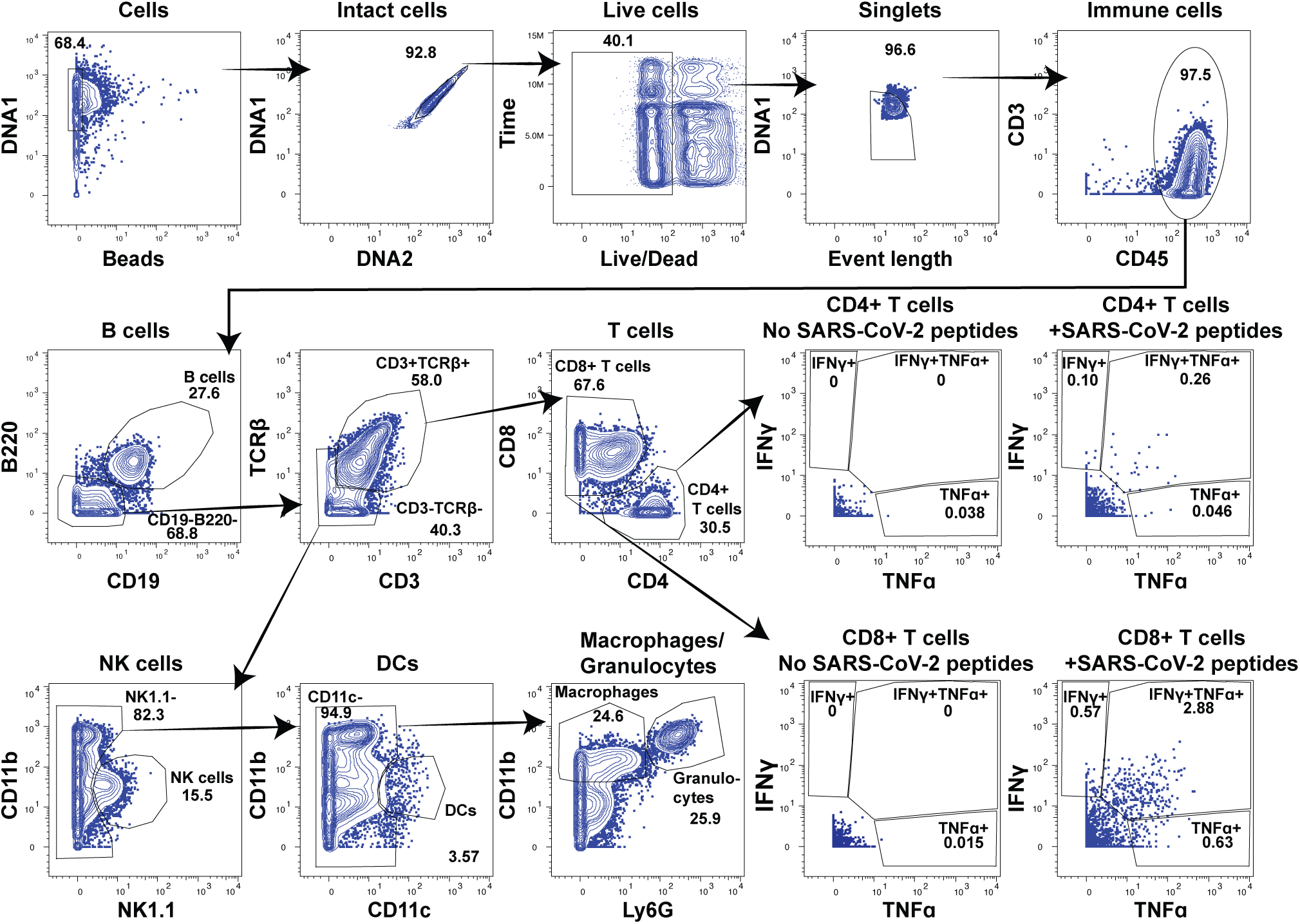
CyTOF gating strategy for identification of pulmonary immune cell subsets and SARS-CoV-2-specific T cells. Shown are the CyTOF gating strategies to identify the indicated immune subsets of cells from the lungs of mice. Datasets were gated on intact, live, singlet cells as indicated, and then for CD45+ cells to identify total immune cells. Gating strategies for B cells, CD4+ T cells, CD8+ T cells, NK cells, DCs, macrophages, and granulocytes are shown. SARS-CoV-2-specific T cells were identified as CD4+ or CD8+ T cells specifically inducing IFNγ and/or TNFα following SARS-CoV-2 peptide stimulation as indicated. Gates are depicted on a representative specimen from the Vac+BA.1 group.

**Figure S2.**
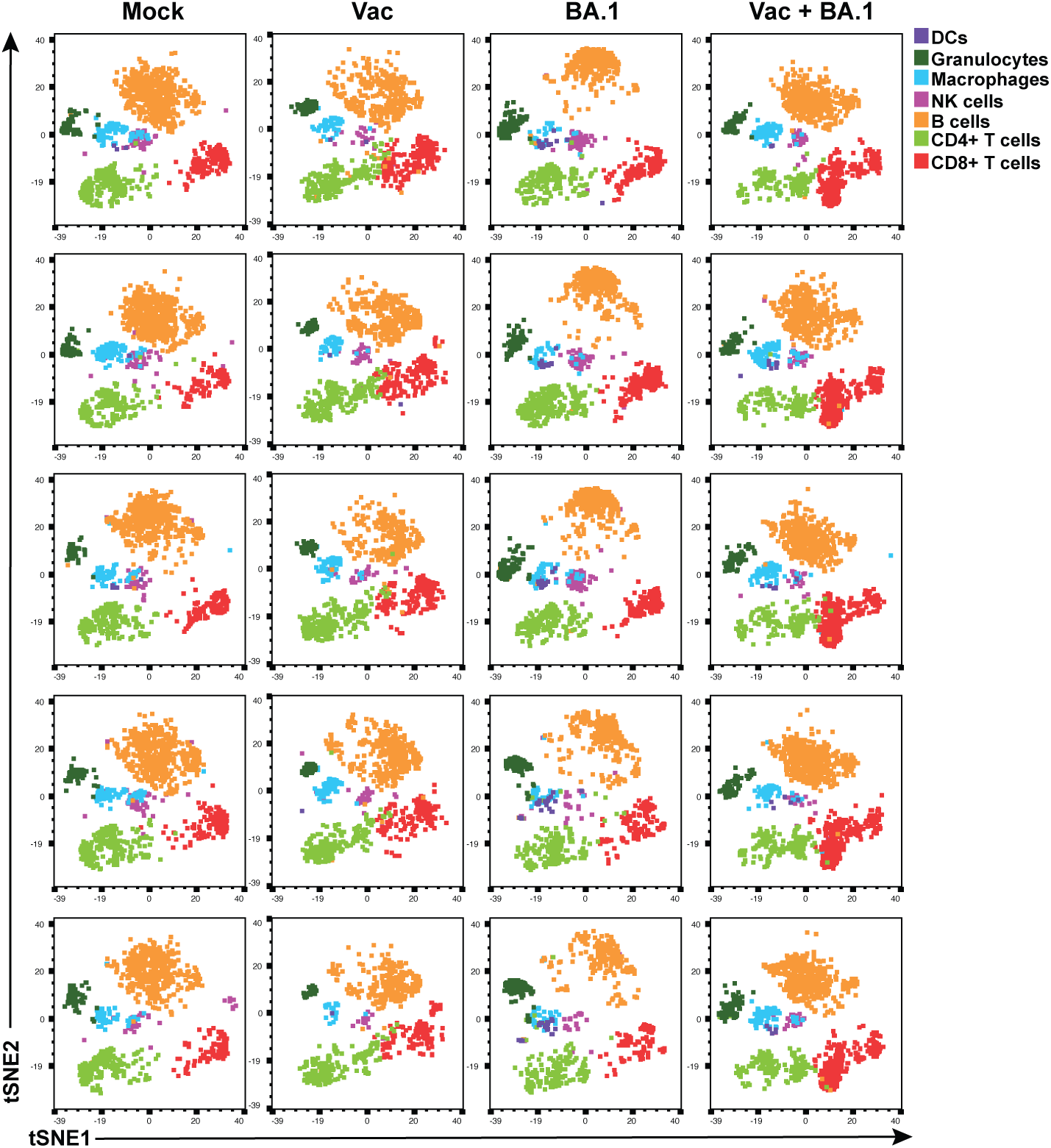
Innate and B cell perturbations persist during BA.1 convalescence. Immune perturbations persist in the lungs of mice 3 weeks after BA.1 infection. The first row depicts data already presented in Fig. 2A, while the remaining rows depict additional mice analyzed in each experimental group. Note overall similarities in cell subset distributions within each group (n=5 mice / group).

**Figure S3.**
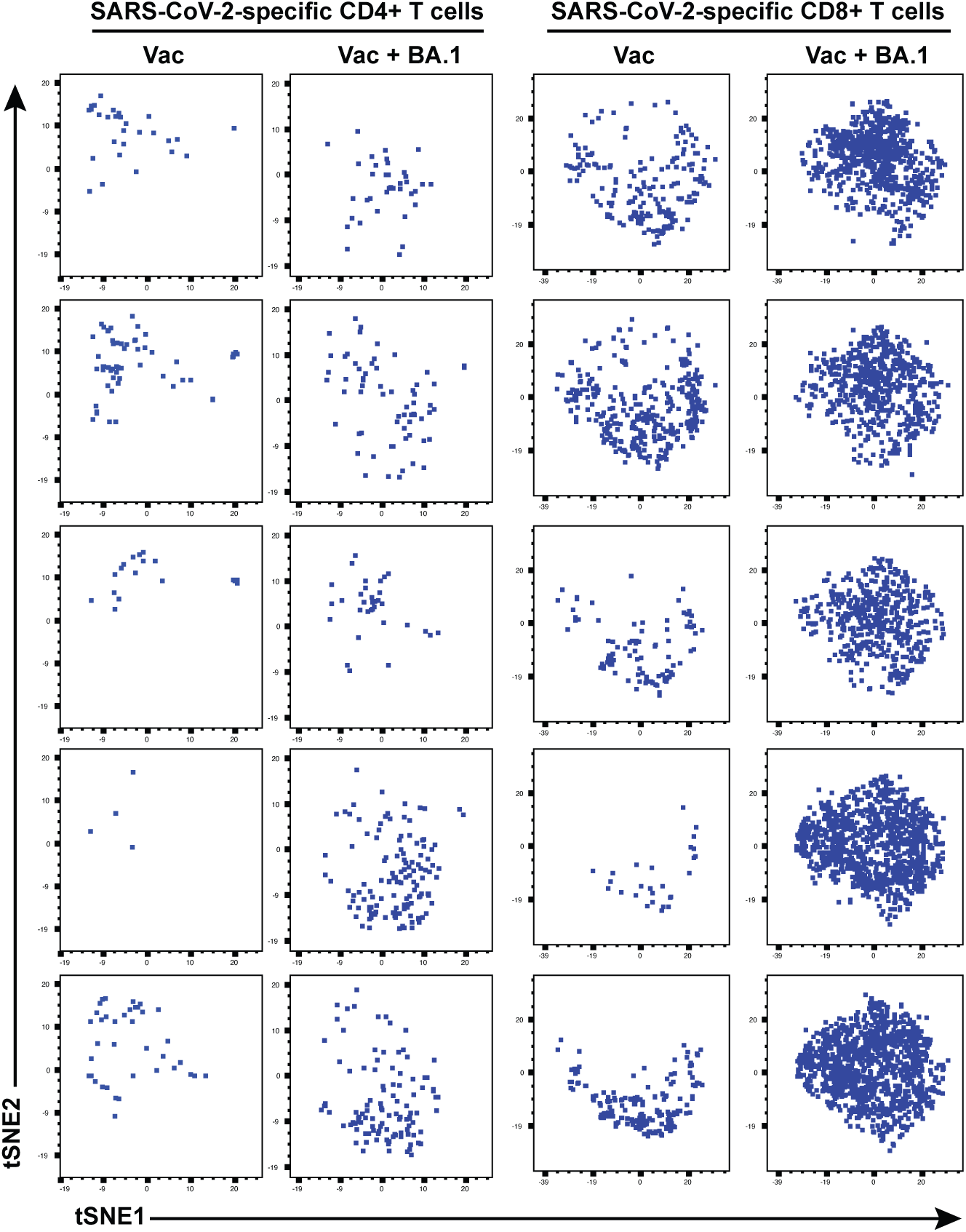
Pulmonary SARS-CoV-2-specific T cells phenotypically differ in Vac as compared to Vac+BA.1 mice. Shown are tSNE plots of pulmonary SARS-CoV-2-specific CD4+ and CD8+ T cells from individual Vac and Vac+BA.1 mice. For each analysis of SARS-CoV-2-specific CD4+ (*left*) and CD8+ (*right*) T cells, each tSNE plot corresponds to a different mouse. The CD4+ T cells were run within one tSNE, while the CD8+ T cells were run within a separate tSNE. The same data concatenated from all 5 mice within each experimental group are shown in Fig. 5A and 5B.

**Figure S4.**
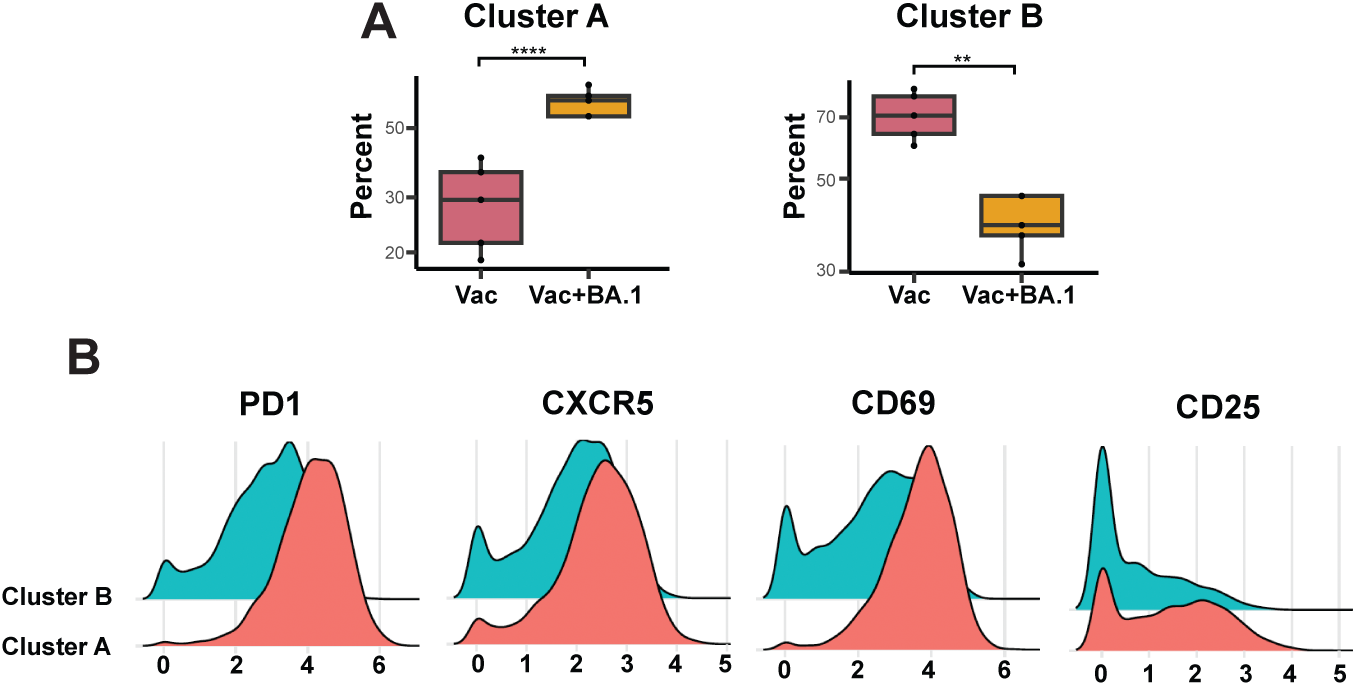
Random Forests-identified cluster features of SARS-CoV-2-specific CD8+ T cells enriched in Vac+BA.1 as compared to Vac mice. (**A**) Unbiased clustering by Random Forests and a silhouette score-based assessment of clustering validity and cross-validation identified 2 clusters of SARS-CoV-2 specific CD8+ T cells. Cluster A was significantly enriched in Vac+BA.1 group, while Cluster B was significantly disenriched in the Vac+BA.1 group, relative to the Vac group. **p<0.01, ****p<0.0001 as assessed using the Welch unpaired t-test and adjusted for multiple testing using the Benjamini-Hochberg for FDR. (**B**) Compared to Cluster B, Cluster A expresses high levels of PD1, CXCR5, CD69, and CD25. Shown are histogram plots corresponding to SARS-CoV-2-specific CD8+ T cells in Clusters A and B.

**Figure S5.**
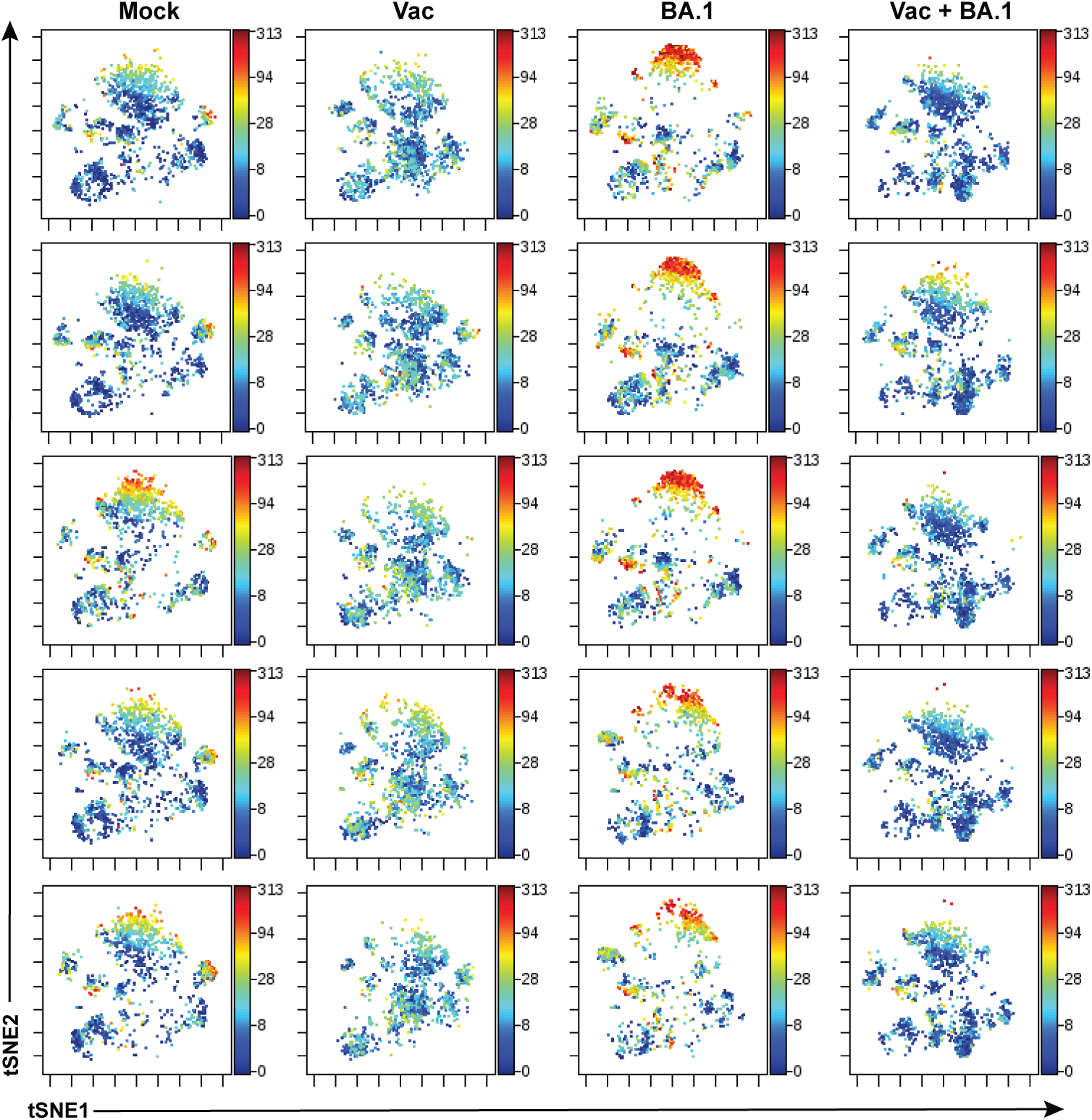
BA.1 convalescent mice, but not BA.1 convalescent mice previously vaccinated, harbor multiple subsets of pulmonary immune cells expressing high levels of CXCR4. The first row depicts data already presented in Fig. 6A, while the remaining rows depict additional mice analyzed in each experimental group. Note preferential high expression levels of CXCR4 among cells of BA.1 convalescent mice.

**Figure S6.**
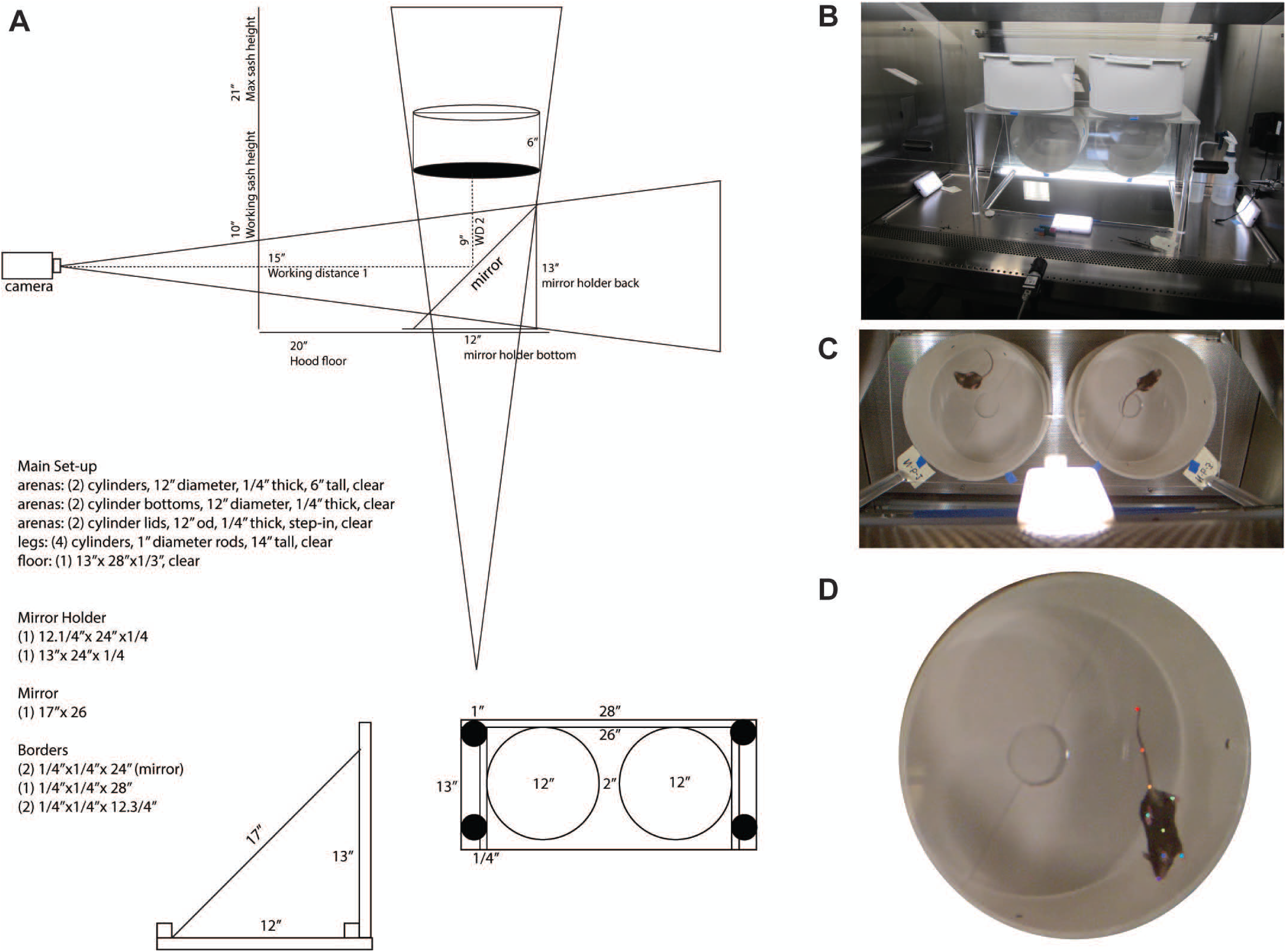
Custom-designed behavioral apparatus for behavioral videography in an ABSL3 biosafety cabinet for behavioral characterization of BA.1 convalescent mice. For simultaneous video acquisition of two independent open field arenas, a mirror was placed at a 45-degree angle to enable ventral views of mouse paws, nose, and tail. Two acrylic cylindrical arenas (12” diameter, 6” tall) with lids made of white corrugated plastic were elevated over the mirror via a clear acrylic platform (13” x 28”). (**A**) Specifications of the behavioral apparatus. (**B**) Photo of behavioral setup configuration in the biosafety cabinet. (**C**) Full camera view of the behavioral arena. (**D**) Behavioral video frame showing DLC labels on tracked body parts.

**Figure S7.**
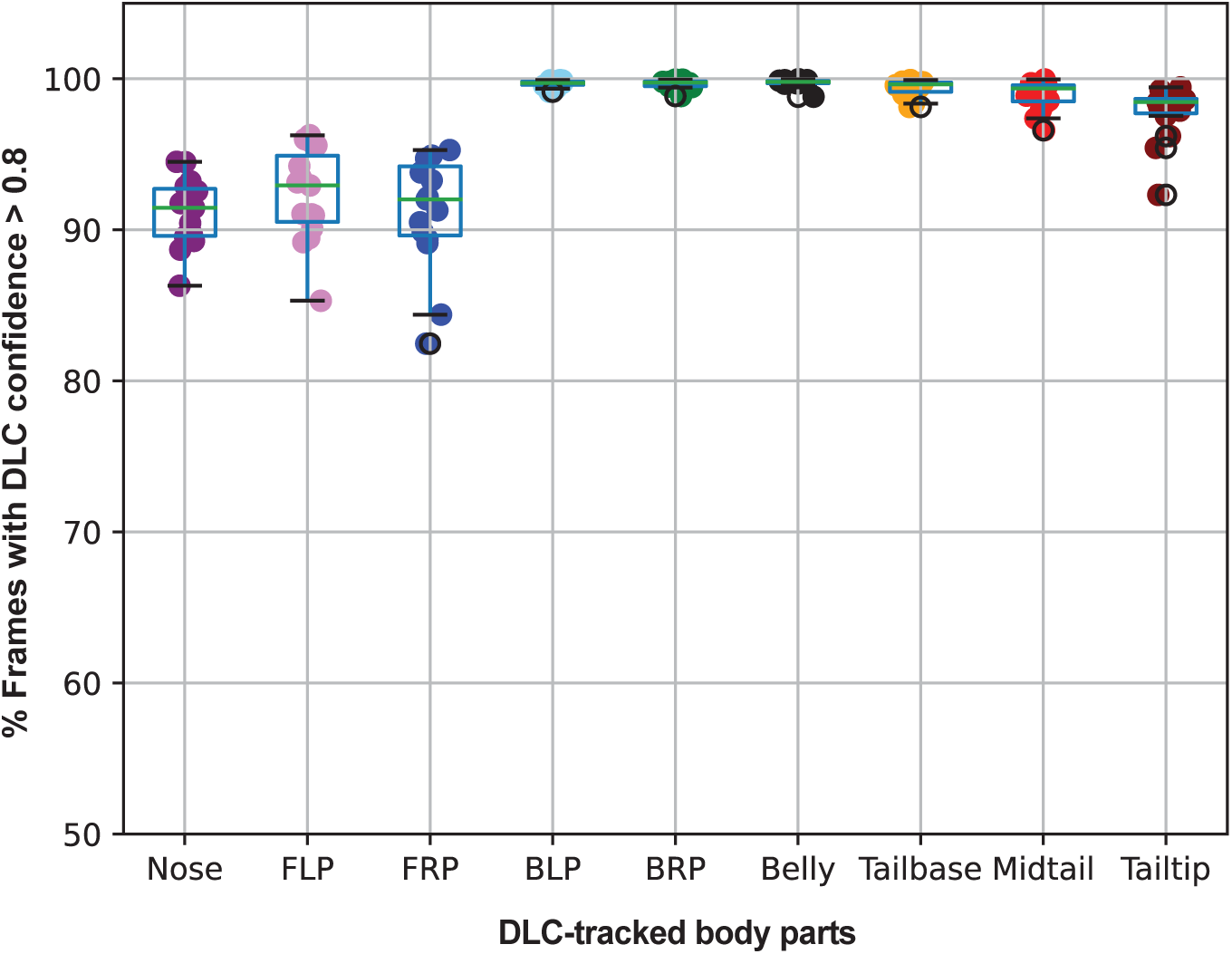
DeepLabCut (DLC) tracking confidence. Percentage of DLC-labeled frames with a confidence value > 0.8 for each body part (colors) over all sessions at DPI 21. Body parts are named as follows: Nose, Front Left Paw (FLP), Front Right Paw (FRP), Back Left Paw (BLP), Back Right Paw (BRP), Belly, Tailbase, Midtail, and Tailtip. Each dot represents one mouse. Lower confidence of the Nose, Front Left Paw, and Front Right Paw are due to occlusions occurring during behaviors that obstruct the camera view of these body parts, such as grooming and rearing.

## ACKNOWLEDGEMENTS

This work was supported by the Van Auken Private Foundation, Roddenberry Foundation, David Henke, Pamela and Edward Taft, James B. Pendleton Charitable Trust, the Gladstone Institutes, and NIH grants RF1AG062234, R01AG062629 and P01AG073082. We also acknowledge the DRC Center Grant P30 DK063720, the S10 1S10OD018040-01 for use of the CyTOF instrument and F31 AI164671. We thank Stanley Tamaki, Claudia Bispo, Vinh Nguyen and Pricsilla Sanchez for CyTOF assistance at the Parnassus Flow Core, Eliver Ghosn for advice on murine lung processing, Françoise Chanut for editorial assistance, and Robin Givens for administrative assistance.

## AUTHOR CONTRIBUTIONS

T.M. designed and performed CyTOF experiments, conducted data analyses, prepared figures and tables, and drafted the manuscript; R.S. designed and performed animal experiments, conducted data analyses, and participated in writing the manuscript. S.R.M. designed animal behavioral experiments, conducted data analyses, and participated in writing the manuscript. K.K.L. performed VAME analysis. R.T. and N.E. performed cluster analysis. K.Y. participated in SPICE analysis, and X.L. established the pipeline for CyTOF data normalization and analysis. N.K. performed animal behavioral data analysis. I.P.C., M.M., and B.S. participated in BSL3 animal experiments. L.S. and J.M. synthesized and purified peptides corresponding to murine epitopes of SARS-CoV-2. F.H.D. provided the mRNA-1273 vaccines. J.J.P., M.O., and N.R.R. conceived the study, performed supervision, and conducted data analyses. N.R.R. wrote the manuscript. All authors have read and approved this manuscript.

